# Role of R5 Pyocin in the Predominance of High-Risk *Pseudomonas aeruginosa* Isolates

**DOI:** 10.1101/2024.10.07.616987

**Authors:** Liyang Zhang, Qi Xu, Filemon C Tan, Yanhan Deng, Morgan Hakki, Samuel A. Shelburne, Natalia V. Kirienko

## Abstract

Infections with antimicrobial resistant pathogens, such as *Pseudomonas aeruginosa,* are a frequent occurrence in healthcare settings. Human *P. aeruginosa* infections are predominantly caused by a small number of sequence types (ST), such as ST235, ST111, and ST175. Although ST111 is recognized as one of the most prevalent high-risk *P. aeruginosa* clones worldwide and frequently exhibits multidrug-resistant or extensively drug-resistant phenotypes, the basis for this dominance remains unclear. In this study, we used a genome-wide transposon insertion library screen to discover that the competitive advantage of ST111 strains over certain non-ST111 strains is through production of R pyocins. We confirmed this finding by showing that competitive dominance was lost by ST111 mutants with R pyocin gene deletions. Further investigation showed that sensitivity to ST111 R pyocin (specifically R5 pyocin) is caused by deficiency in the O-antigen ligase *waaL*, which leaves lipopolysaccharide (LPS) bereft of O antigen, enabling pyocins to bind the LPS core. In contrast, sensitivity of *waaL* mutants to R1 or R2 pyocins depended on additional genomic changes. In addition, we found the PA14 mutants in lipopolysaccharide biosynthesis (*waaL*, *wbpL*, *wbpM*) that cause high susceptibility to R pyocins also exhibit poor swimming motility. Analysis of 5,135 typed *P. aeruginosa* strains revealed that several international, high-risk sequence types (including ST235, ST111, and ST175) are enriched for R5 pyocin production, indicating a correlation between these phenotypes and suggesting a novel approach for evaluating risk from emerging prevalent *P. aeruginosa* strains. Overall, our study sheds light on the mechanisms underlying the dominance of ST111 strains and highlighting the role of *waaL* in extending spectrum of R pyocin susceptibility.

## Introduction

The inhabitants of every biological niche strive to obtain as much space and as many resources as possible, engaging in dynamic processes of competition and cooperation. For pathogenic bacteria, the human body is an excellent biological niche, and innumerable such interactions occur constantly. Interactions between the Gram-negative bacterium *Pseudomonas aeruginosa* and other pathogens have been particularly thoroughly studied (1–5). *P. aeruginosa* has several advantages over other bacteria, including the production of robust biofilms (6, 7), the secretion of toxic factors into the extracellular environment or directly into target cells (8–11), and innate resistance to multiple antibiotics (12, 13), all of which support efficient host colonization in the healthcare setting.

Interactions between strains of *P. aeruginosa* are also complex, with some strains inhibiting other’s growth (14–16). Prior work from several labs, including ours, revealed that the multilocus sequence type (MLST) group ST111 dominates in hematopoietic cell transplant (HCT) recipients and hematologic malignancy (HM) patients with bloodstream infections (BSI) at Oregon Health and Science University (OHSU), accounting for 38.6% (22/57) of isolates (17, 18). *P. aeruginosa* ST111 has been documented as one of several high-risk, multidrug- or extensively drug resistant clones that are prevalent worldwide (18–22). We determined that ST111 isolates from these patients out-competed non-ST111 isolates via live-cell-dependent and live-cell-independent mechanisms (17). However, the specific mechanisms underlying these observations and the relative importance of these mechanisms remain poorly defined.

One of the key mechanisms used for intrastrain competition amongst *P. aeruginosa* strains is the production of high-molecular weight bacteriocins, called pyocins, that are categorized into three groups: R, S, and F (14, 23). R pyocins, the most common class, are phage-tail-like complexes related to the P2 bacteriophage (24). R pyocin tail fibers attach to lipopolysaccharide (LPS) on the cell surface of recipient cells, the sheath contracts, and the core structure is injected through the cell membrane (23, 25–30). This depolarizes the cell membrane and prevents active amino acid transport (31). The presence of the O-antigen on the outermost region of LPS limits its interaction with R pyocins and confers protection (26). Based on amino acid sequence variations and their ability to kill competing bacteria, R pyocins are further divided into R1-R5 subtypes (23). Of these, R5 is thought to have the broadest activity, with a bactericidal spectrum that includes the ranges of all the other subtypes (32, 33). The importance of R pyocins in interstrain and interspecies competition has been previously studied (14, 34–36). To date, however, little has been reported on the roles of R pyocin subtypes in *P. aeruginosa* strain dominance and its impact in clinical settings.

Using an unbiased approach, this study identified an important role for R5 pyocins in the predominance of *P. aeruginosa* ST111. We found that ST111 isolates inhibited a large panel of non-ST111 strains by producing a high-molecular weight bactericidal product. A genome-wide, transposon-insertion mutant screen identified this product as R pyocin. We also determined that loss of function of the O-antigen ligase WaaL caused the susceptibility of previously resistant strains to R pyocins. Interestingly, lipopolysaccharide biosynthesis mutants with increased susceptibility to R5 pyocins (such as PA14*ΔwaaL*) also had reduced swimming motility, suggesting that this difference may provide a useful diagnostic tool. Deletion of an R5 pyocin structural gene abolished ST111’s advantage in intraspecies competition, establishing its importance. This is also consistent with results seen from large-scale data mining of *P. aeruginosa* isolate genomes worldwide, which demonstrated a strong correlation between the presence of R5 pyocin and the prevalence of multiple high-risk clones, including ST235, the most prevalent sequence type across the globe.

## Results

### An extracellular bactericidal complex contributes to the competitive advantage of ST111 strains

Previously, we and others described *P. aeruginosa* sequence type ST111 as predominant among clinical isolates from HCT/HM BSI patients at OHSU (17, 18). Almost all the ST111 isolates possessed inactivating mutations in the carbapenem entry porin OprD. While our previous work and that of others demonstrated a competitive advantage for OprD inactivation (17, 37), this factor was unlikely to explain the dominance of ST111 alone; most of the non-ST111 clinical isolates also carried loss-of-function mutations in *oprD*. For this reason, we investigated other possible mechanisms for the dominance of ST111 isolates over clinical isolates from other sequence types.

To observe the interactions between ST111 and non-ST111 isolates, we first tested *in vitro* competition between strains with *oprD* mutations: a carbenicillin-susceptible ST111 strain, M0101 (has *oprD* mutation), and two *oprD-*mutant, non-ST111 strains (ST291/M0103::Carb^R^ and ST233/M0104::Carb^R^). Immediately after mixing, ST291/M0103::Carb^R^ or ST233/M0104::Carb^R^ comprised approximately half of the co-culture (Fig 1A). After 24 hours co-incubation in the absence of antimicrobials, we observed that carbenicillin-resistant, non-ST111 bacteria represented only a small fraction of the overall population, especially when compared to the initial input (Fig 1A). This was not due to the growth deficiencies or loss of the carbenicillin resistance plasmid: ST291/M0103::Carb^R^ and ST233/M0104::Carb^R^ grew well on carbenicillin after 24 h incubation in antibiotic-free medium in the absence of ST111 strain (Fig 1A). Similar results were seen in a *Caenorhabditis elegans*-based, *in vivo* competition assay, where ST111/M0101 prevented ST291/M0103::Carb^R^ or ST233/M0104::Carb^R^ cells from colonizing the nematode host (Fig 1B).

**Figure 1.**
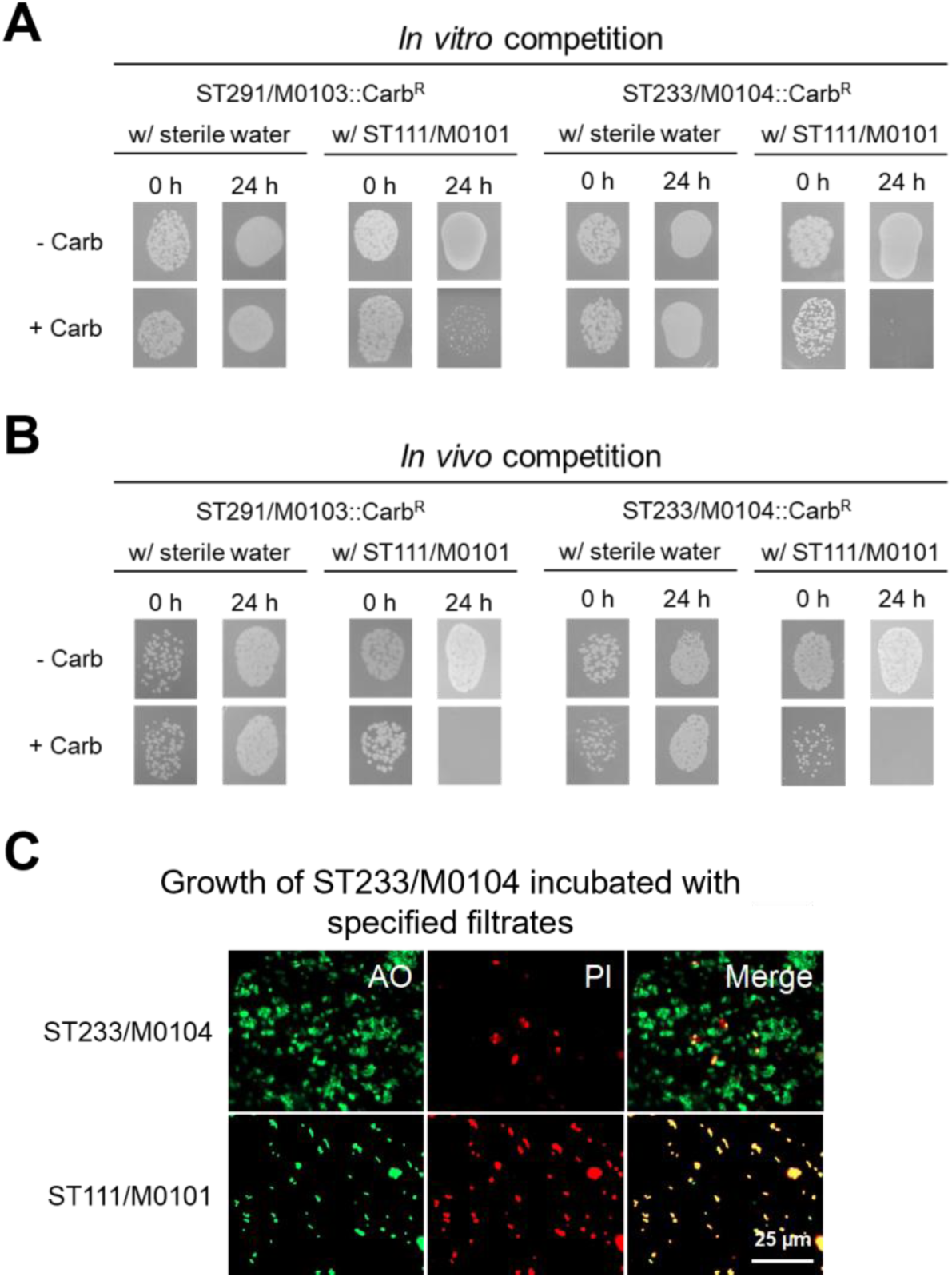
ST111 strains outcompete non-ST111 clinical isolates. (**A, B**) Bacterial colonies of ST111/M0101 and carbenicillin-resistant non-ST111 (ST291/M0103::Carb^R^ or ST233/M0104::Carb^R^) strains on non-selective LB plates (-Carb) and carbenicillin-containing LB plates (+ Carb) before (0 hours) and after (24 hours) an *in vitro* competition (**A**) or an *in vivo* competition in *C. elegans* (**B**). In conditions without antimicrobial, both bacterial strains grow; in the presence of carbenicillin, there is selection only for the non-ST111 strain. Three biological replicates were performed with three technical replicates for each. (**C**) Images of ST233/M0104 cells stained with acridine orange (AO, a cell-permeant dye) and propidium iodide (PI, a cell-impermeant dye that stains only dead cells) after 4 hours of incubation with self *vs.* ST111/M0101 filtrate. Three biological replicates were performed with three technical replicates for each. Scale bar: 25 μm. (**A-C**) A representative technical replicate is shown for each condition.

We postulated that ST111 strains may be outcompeting non-ST111 strains by releasing a toxic (bactericidal or bacteriostatic) factor. Supporting this idea, cell-free spent media (hereafter referred to as filtrate) from ST111 strains (ST111/M0067, ST111/M0101, or ST111/218M0087) impaired the growth of several non-ST111 strains (ST233/M0104, ST291/M0103, or ST299/M0128) (Figs S1A-C, also see Table 1). Interestingly, this growth-suppressing ability was specific. Neither ST111 nor ST446 strains showed apparent growth inhibition in the presence of ST111 filtrate (Table 1). ST446 is the second-most frequent sequence type amongst *P. aeruginosa* bloodstream infection isolates collected from OHSU (17). We also observed that ST446 filtrates inhibited certain non-ST111 strains (Fig S1D-F). Follow-up experiments showed that the factor of interest, which we presumed was a toxin, is temperature-sensitive and is retained after centrifugation through 100-kDa membrane (Figs S2A-C). Interestingly, filtrate from the laboratory-adapted strain ST253/PA14 also inhibited the same three non-ST111 strains (ST291/M0103, ST233/M0104, or ST299/M0128) (Figs S2D-F). As with ST111 filtrate, growth inhibition from this filtrate was lost after heat treatment or removal of macromolecules >100 kDa (Figs S2D-F).

**Table 1.** Growth inhibition by specified filtrates.

| Inhibit the growth of | MLST | filtrate from |  |  |  |  |
| --- | --- | --- | --- | --- | --- | --- |
|  |  | PA14 253 | PAO1 549 | M0067 111 | M0101 111 | 218M0087 111 |
| M0067 | 111 | – | – | – | – | – |
| M0101 | 111 | – | – | – | – | – |
| 218M0087 | 111 | – | – | – | – | – |
| M0003 | 111 | – | – | – | – | – |
| M0025 | 111 | – | – | – | – | – |
| M0134 | 111 | – | – | – | – | – |
| M0169 | 111 | – | – | – | – | – |
| M0177 | 111 | – | – | – | – | – |
| M0249 | 111 | – | – | – | – | – |
| M0117 | 446 | – | – | – | – | – |
| M0186 | 446 | – | – | – | – | – |
| M0068 | 132 | + | + | + | + | + |
| M0103 | 291 | + | + | + | + | + |
| M0104 | 233 | + | + | + | + | + |
| M0128 | 299 | + | + | + | + | + |
| M0013 | 17 | + | + | + | + | + |
| M0027 | 281 | + | + | + | + | + |
| M0043 | 308 | + | + | + | + | + |
| M0089 | 260 | + | + | + | + | + |

MLST: Multilocus sequence typing; –: not inhibited; +: inhibited

To distinguish whether ST111 filtrates were bactericidal or bacteriostatic against non-ST111 strains, we visualized the interaction between filtrates and strains by nucleic acid staining with cell-permeant acridine orange and cell-impermeant propidium iodide dyes. After four hours of incubation with ST111/M0101 filtrate, the fraction of ST233/M0104 cells labeled with propidium iodide compared to those treated with self-filtrate control significantly increased, indicating that the ST111 or PA14 filtrates caused cell permeabilization and death (Fig 1C and Fig S2G).

### High-throughput genetic screening identifies R pyocin as the growth inhibitor

Based on these experimental outcomes, the most parsimonious hypothesis was that ST253/PA14 and ST111 produced similar, if not the same, toxic, high-molecular-weight complex. To identify any factor(s) responsible for bactericidal activity in ST253/PA14 and ST111 filtrates, we performed a high-throughput, genetic screen using a commercially-available, non-redundant PA14 transposon mutant library (38). Instead of producing individual filtrates from each of the thousands of clones, the growth of overnight cultures of ST253/PA14 transposon mutants were suppressed with meropenem, initially for 5 h at 16 µg/mL and then for an additional ∼ 18 h at 8 µg/mL after mixing with the non-ST111, meropenem-resistant sensor strain ST233/M0104 (Fig 2A). To act as a “sensor strain” for growth inhibition, ST233/M0104 was engineered to constitutively expresses dsRed, enabling bacterial growth to be assessed via fluorescence measurement (Fig 2A). The assay was validated using wild-type ST253/PA14, *Escherichia coli* OP50, and *Enterococcus faecalis* OG1RF (Fig S3). 5,459 transposon insertion mutants were screened, covering ∼90% of ST253/PA14 protein-coding genes. 43 hits (0.79% hit rate) were identified in the primary screen (Fig 2B) based on increased bacterial growth, as assessed by fluorescence (>3 standard deviations above the mean) (Fig 2C).

**Figure 2.**
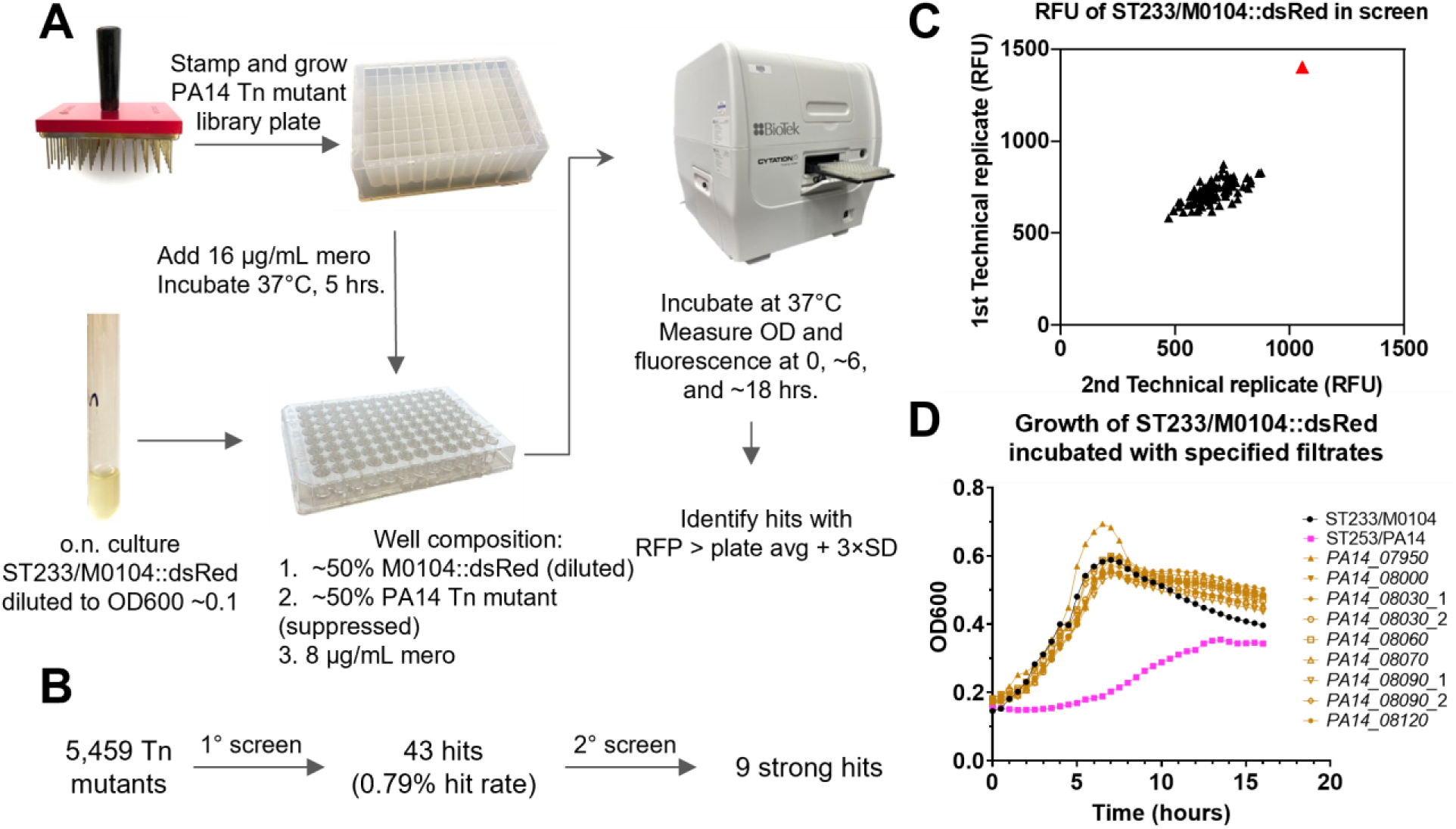
Genome-wide transposon screening identifies R pyocin as the relevant bactericidal agent. **(A)** Schematic of primary screen. Two technical replicates were performed for each library plate. Tn: Transposon insertion; o.n.: overnight; mero: meropenem; avg: average; SD: standard deviation. **(B)** Summary of screen results. **(C)** Representative primary screen data from 96-well library plate. M0104::dsRed growth was measured via red fluorescence after 18 hours. Read from the well containing the verified hit *PA14_08030*_1 is highlighted in red. (**D**) Representative secondary screen ST233/M0104 growth curves in the presence of 50% filtrate from self ST233/M0104 (negative control), WT ST253/PA14 (positive control), or 9 sequence-confirmed ST253/PA14 strong mutant hits, which are related to pyocin (gold). Three biological replicates with two technical replicates each were performed for all primary screen hits.

While the use of antibiotic-treated cultures made the primary screen feasible, treatment-mediated bacterial lysis could cause the release of intracellular factors not normally present in the filtrate, altering the growth of the sensor strain. Therefore, we validated our primary hits using filtrates against the sensor strain ST233/M0104 (Fig 2D). After testing in two additional sensitive strains, ST291/M0103::dsRed, or ST299/M0128::dsRed, 9 strong hits emerged (Table 2 and Fig S4). None of these mutants showed significant growth defects, confirming that the growth inhibition observed was a specific phenomenon (Fig S5A).

**Table 2.** List of 9 strong hits.

| Gene Name | Gene annotation | Sensor Strain Growth |  |  |
| --- | --- | --- | --- | --- |
|  |  | ST233/M0104 | ST291/M0103 | ST299/M0128 |
| *PA14_07950/prtN* | [pyocin] transcriptional regulator | Strong | Strong | Strong |
| *PA14_08000* | conserved hypothetical [pyocin collar] protein | Strong | Strong | Strong |
| *PA14_08030_1* | putative phage baseplate assembly protein | Strong | Strong | Strong |
| *PA14_08030_2* | putative phage baseplate assembly protein | Strong | Strong | Strong |
| *PA14_08060* | putative tail fiber assembly protein | Strong | Strong | Strong |
| *PA14_08070* | phage tail sheath protein | Strong | Strong | Strong |
| *PA14_08090_1* | putative phage tail tube protein | Strong | Strong | Strong |
| *PA14_08090_2* | putative phage tail tube protein | Strong | Strong | Strong |
| *PA14_08120* | putative tail length determinator protein | Strong | Strong | Strong |

Eight of the transposon mutants that abolished PA14-mediated growth inhibition of ST233/M0104 were in genes that encoded structural components of R pyocin (Table 2), a bacteriocin produced by *P. aeruginosa* (23). R pyocins resemble the contractile tail structure of the P2 bacteriophages and can disrupt cell membranes by penetration (26, 27, 29, 30). Identification of a role for R pyocin was consistent with our observations that filtrate toxicity was associated with a high-molecular-weight, bactericidal effector. The ninth strong hit was a transposon insertion in the positive transcriptional regulator *prtN,* which drives expression of the R pyocin structural genes (39, 40). Mutations in *prtN* have been shown to abolish R pyocin production in *P. aeruginosa* (40).

To measure R pyocin production in our 9 hits, filtrates were tested for their ability to inhibit growth of *P. aeruginosa* 13s, an indicator strain for R pyocins (41). As expected, filtrates from mutants of ST253/PA14 with insertions in the R pyocin genes did not inhibit the growth of *P. aeruginosa* 13s. Likewise, a PA14*Δpyocins* mutant, where the operons for the R and F pyocins were deleted (42), failed to inhibit growth of the 13s indicator strain. Taken together, these data support the conclusion that R pyocin production was compromised in our transposon mutants (Fig S5B). Combined, these data strongly argue that R pyocins are the primary mechanism driving ST253/PA14-mediated inhibition of the growth of non-ST111 clinical isolates.

### R pyocin is required for ST111 dominance against non-ST111 strains

Genome sequence analysis of OHSU ST111 strains revealed that they all encoded R pyocins. Filtrates from each of the three ST111 strains (ST111/M0067, ST111/M0101, or ST111/218M0087) strongly inhibited growth of the R pyocin indicator strain 13s (Fig 3A), confirming production of R pyocins.

**Figure 3.**
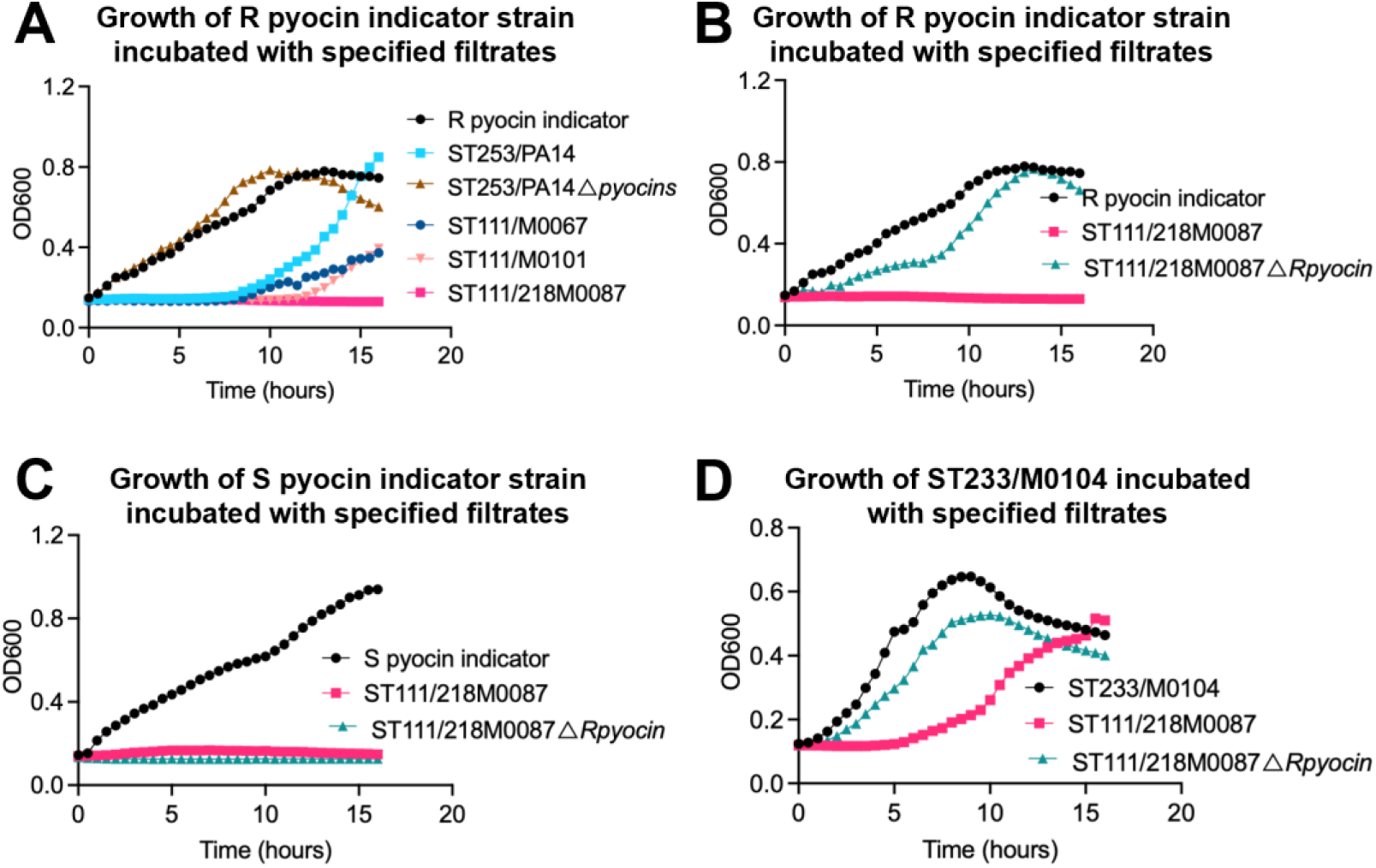
R pyocin mediates the growth inhibition exhibited by ST111 isolate filtrates. (**A**) Growth of the R pyocin indicator strain *P. aeruginosa* 13s incubated with filtrates from self, ST253/PA14, ST253/PA14*Δpyocins*, ST111/M0067, ST111/M0101, or ST111/218M0087. (**B-D**) Growth of the R pyocin indicator strain 13s (**B**), S pyocin indicator strain PML1516d (**C**), or ST233/M0104 (**D**) incubated with filtrates from wild-type ST111/218M0087 or ST111/218M0087*ΔRpyocin* (in this strain, an ortholog of structural pyocin gene (tail tube) *PA14_08090* was deleted). Three biological replicates with three technical replicates each were performed.

As pyocins come in three major types, S, F, and R (23), we tested our panel of ST111 isolates for S pyocins using the indicator strain PML1516d. As anticipated, the three strains produced S type pyocins as well (Fig S6A). To determine whether S pyocins contributed to the inhibition of non-ST111 isolates, we deleted the putative R pyocin tail tube protein gene in ST111/218M0087. This gene was selected based on its sequence homology to *PA14_08090*, which encodes phage tail tube protein in PA14. As expected, removal of the gene strongly compromised the ability of ST111/218M0087*ΔRpyocin* to inhibit the growth of the R pyocin indicator strain (Fig 3B). Loss of this gene did not affect the ability of ST111/218M0087*ΔRpyocin* to prevent growth of the S pyocin indicator strain (Fig 3C). Importantly, ST111/218M0087*ΔRpyocin* lost its inhibitory activity against the non-ST111 panel (Fig 3D and Fig S6B-C).

To more quantitatively assess the impact of R pyocin deletion on toxicity, we counted colony-forming units (CFUs) to analyze cell viability after filtrate treatment. After one hour of exposure to ST111/218M0087 filtrate, there were no viable ST260/M0089, ST291/M0103, or ST299/M0128 cells, while the number of viable bacteria after exposure to filtrate from the deletion mutant, ST111/218M0087*Δrpyocin*, were comparable to their own filtrates (Fig 4A), demonstrating rapid and strong bactericidal activity of R pyocins. These data indicate that R, but not S, pyocins play a crucial role in filtrate-dependent killing of sensitive strains. Similarly, bactericidal properties of ST253/PA14 filtrate were abolished in filtrate from an analogous ST253/PA14*Δpyocins* deletion mutant (Fig S6D).

**Figure 4.**
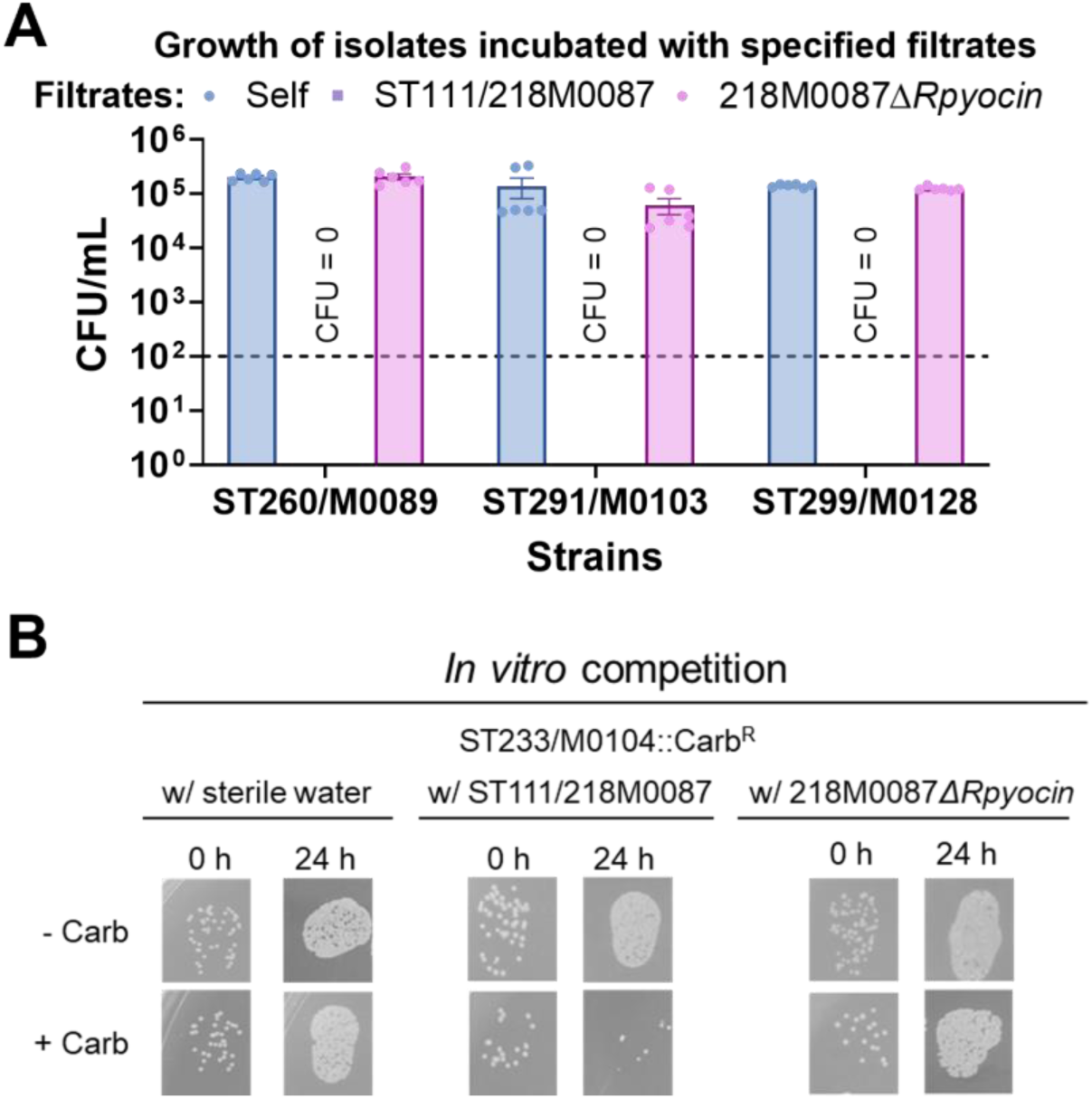
R pyocin deletion causes loss of ST111 dominance. CFUs (colony-forming units) of ST260/M0089, ST291/M0103, and ST299/M0128 after 1 hour incubation with self, ST111/218M0087, and ST111/218M0087*ΔRpyocin* filtrates, respectively. Three biological replicates were performed. Detection limit of CFU assay (horizontal dashed line) is 100 colonies/mL per individual technical replicate. Error bars represent SEM. (**B**) Bacterial colonies of ST111/218M0087, or ST111/218M0087*ΔRpyocin* and carbenicillin-resistant non-ST111 (ST233/M0104::Carb^R^) strains on non-selective LB plates (-Carb) and carbenicillin-containing LB plates (+ Carb) before (0 hours) and after (24 hours) an *in vitro* competition. In conditions without antimicrobial, both bacterial strains grow; in the presence of carbenicillin, there is selection only for the non-ST111 strain. Three biological replicates were performed with three technical replicates for each. For (**B**), a representative technical replicate is shown for each condition.

R pyocin complexes resemble phage tails, and *P. aeruginosa* is known to produce endogenous phages. To explore the role of phages in filtrate-dependent killing of non-ST111 clinical isolates (43), we spread filtrate from ST233/M0104, ST253/PA14, ST111/218M0087, or ST111/218M0087*ΔRpyocin* on a confluent lawn of ST233/M0104 bacteria. We noted that ST233/M0104 is an endogenous phage carrier. However, no notable increase in plaques was observed following treatment with filtrates from ST253/PA14, ST111/218M0087, or ST111/218M0087*ΔRpyocin* (Fig S7). Notably, the addition of 200 μL of filtrate from ST253/PA14 or ST111/218M0087 to a lawn of ST233/M0104 resulted in significant bacterial killing, a phenomenon distinctly different from the effects seen with the filtrate from ST233/M0104 or ST111/218M0087*ΔRpyocin* (Fig S7B). Combined, these data indicate that the growth inhibition is specific to the production of R pyocins and is independent of the presence of phages.

Importantly, compromising R pyocin production also limited growth inhibition of non-ST111 strains during *in vitro* competition assays. Co-incubation (24 h) of wild-type ST111/218M0087 with ST233/M0104::Carb^R^ strongly inhibited growth of the latter, while the 24-hour co-incubation of ST111/218M0087*ΔRpyocin* with ST233/M0104::Carb^R^ led to titers in the latter comparable to single-strain controls (Fig 4B).

### Sensitivity towards ST111 R pyocins correlates with swimming motility

Several studies have demonstrated that R pyocins target the core of LPS on the outer membrane of *P. aeruginosa* (44, 45). This interaction is hindered by the addition of O-antigens onto LPS (26). We hypothesized that non-ST111 strains may be susceptible to R pyocins due to a deficiency in LPS biosynthesis. To test this, a panel of LPS mutants from the ST253/PA14 transposon mutant library and a PA14*ΔwaaL* mutant we generated were tested for R pyocin susceptibility. Of the eighteen mutants tested, three (waaL, wbpL, and wbpM) showed an increase in sensitivity to ST111 filtrate (Table 3), suggesting that these genes encode proteins that promote resistance. This is consistent with a previous report that *wbpL* and *wbpM* mutations increased susceptibility to R1 and R5 pyocins (26). Therefore we hypothesized that WaaL, the O-antigen ligase mediating the addition of the O-antigen to the LPS core, also plays a key role in determining sensitivity to R pyocin. Despite this, the *PA14ΔwaaL* mutant was not affected by the R pyocin from ST253/PA14 (Table 3).

**Table 3.** R pyocin sensitivity of PA14 LPS mutants.

|  | Gene Annotation | ST253/PA14<br>filtrate | ST111/218M0087<br>filtrate |
| --- | --- | --- | --- |
| *galU* | UTP-glucose-1-phosphate<br>uridylyltransferase | — | — |
| *gmd* | GDP-mannose 4,6-dehydratase | — | — |
| *lptD* | LPS-assembly protein | — | — |
| *lpxC* | UDP-3-O-[3-hydroxymyristoyl] N-acetylglucosamine deacetylase | — | — |
| *lpxO1* | lipopolysaccharide biosynthetic protein | — | — |
| *lpxO2* | lipopolysaccharide biosynthetic protein | — | — |
| *PA14_69170* | probable O-antigen acetylase | — | — |
| *pagL* | Lipid A 3-O-deacylase | — | — |
| *waaL* | O-antigen ligase | — | + |
| *wbpL (orfN)* | glycosyltransferase | — | + |
| *wbpM* | nucleotide sugar epimerase/dehydratase | — | + |
| *wbpW* | phosphomannose isomerase/GDP-mannose | — | — |
| *wbpX* | glycosyltransferase | — | — |
| *wbpY* | glycosyltransferase | — | — |
| *wbpZ* | glycosyltransferase | — | — |
| *wzm* | membrane subunit of A-band LPS efflux<br>transporter | — | — |
| *wzt* | ABC subunit of A-band LPS efflux<br>transporter | — | — |
| *wzz* | O-antigen chain length regulator | — | — |

—: not inhibited; +: inhibited

Previous work from Abeyrathne et al. revealed that *waaL* mutation reduces swimming motility (46). Swimming motility is a form of movement in *P. aeruginosa* that requires a functioning polar flagellum (47). Mutants without a functional flagellum exhibited limited swimming motility. For example, deletion of the flagellar hook-associated protein FlgK (PA14*ΔflgK*) strongly compromised swimming and was used as a positive control (Fig 5A). We hypothesized that swimming motility is related to LPS biosynthesis and that, on this basis, swimming motility may serve as a useful mechanism for predicting susceptibility to R pyocins. To test this prediction, we used a plate-based swimming motility assay to measure swimming motility in wild-type, PA14*ΔflgK*, and PA14 LPS mutants. For each strain, the size of the colony, representing the distance traveled by swimming bacteria and referred to as the swimming diameter (46, 47), was measured. As predicted, reduced swimming motility was observed in the three mutants that caused susceptibility to ST111 R pyocins (*waaL*, *wbpL*, and *wbpM*) (Figs 5A, 5B, and Fig S8A).

**Figure 5.**
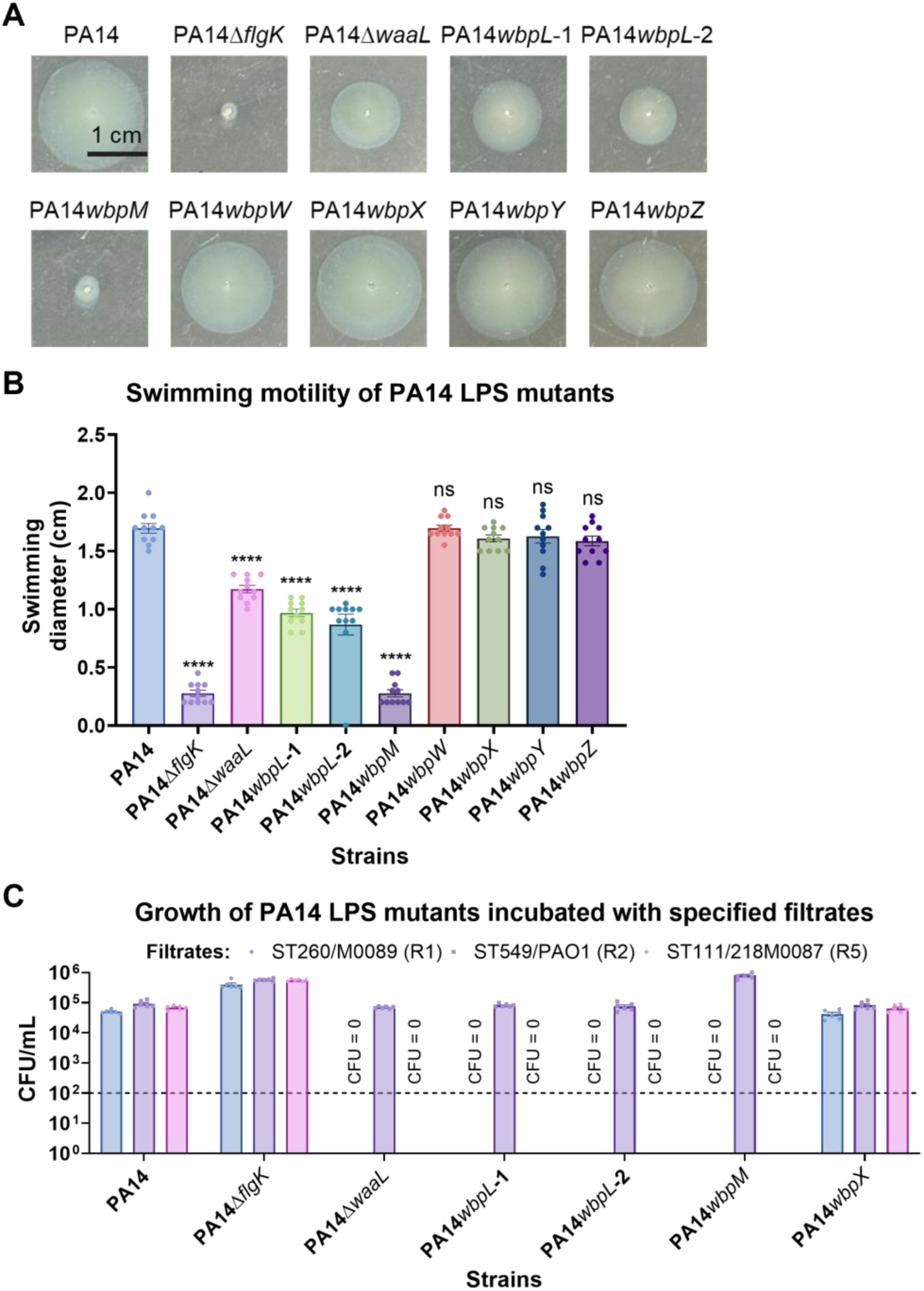
Deficiency in LPS biosynthesis is associated with the high susceptibility to R pyocins. **(A)** The cropped pictures showing the swimming motility of *P. aeruginosa* PA14 mutants. A representative technical replicate is shown for each strain. The original pictures are included in Fig S8A. (**B**) The swimming diameter of WT PA14 and isogenic mutants in the plate-based swimming motility assay. Four biological replicates were performed with at least two technical replicates for each. (**C**) CFUs (colony-forming units) of PA14 mutants after 1 hour incubation with filtrates from an R1 pyocin-encoding strain ST260/M0089, a R2 pyocin-encoding strain ST549/PAO1, or a R5 pyocin-encoding strain ST111/218M0087 respectively. Three biological replicates were performed. Detection limit of CFU assay (dashed horizontal line) is 100 colonies/mL per individual technical replicate.

### Pyocin sensitivity of *waaL* mutants depends on R pyocin type and the presence of self pyocin

Previous research has established that different R pyocin subtypes exhibit differences in bactericidal properties (26). We anticipate that this is due to structural differences, so we compared the sequences of R pyocin tail fiber proteins of our clinical isolates to ST253/PA14 or ST549/PAO1 and found several polymorphisms unique to ST111 strains (Table 4), most of which were located on the tail fiber protein (*PA14_08050* ortholog), with five of eight tail fiber mutations in the Knob1 domain (28). Interestingly, we noticed that an isolate from the ST446 sequence type, the second-most dominant sequence type amongst our clinical isolates, shared the ST446 tail fiber polymorphism. R pyocins produced by *P. aeruginosa* have been placed in five groups based on their bactericidal activity (48). R pyocins from ST111 isolates belong to the R5 subtype, which has the broadest killing spectrum and may contribute to the fitness advantage of this dominant sequence type (32). ST253/PA14 and PAO1 both encode R2 pyocins. Sequence similarity between tail fiber proteins in the R2, R3, and R4 pyocin subgroups is over 98%, so we treated them as a single category (referred to as the R2 group).

**Table 4.**
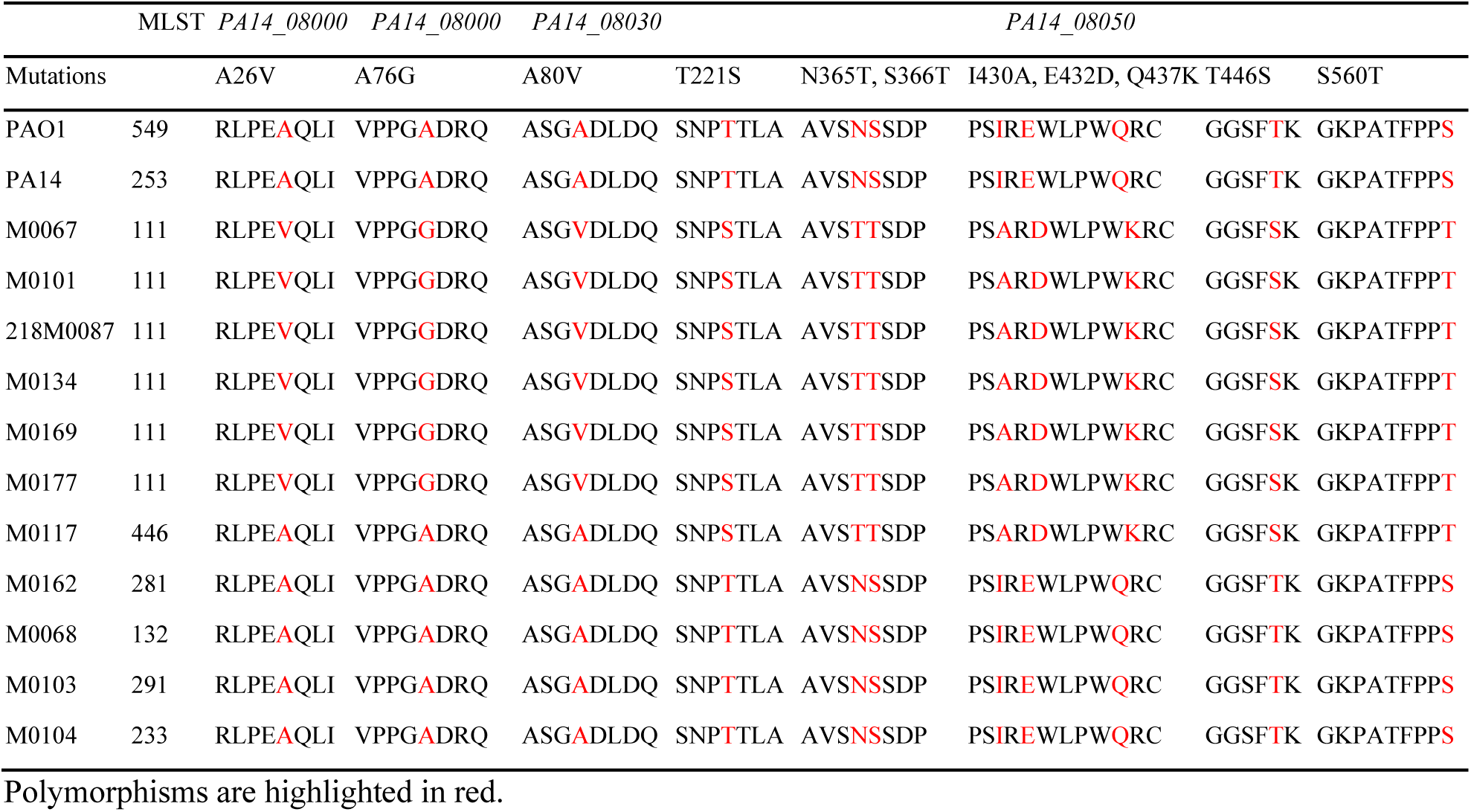
Sequence polymorphisms in different R-type pyocins.

Sequence analysis indicates that R1 pyocin shares a higher degree of homology with R5 pyocin than with R2 pyocin, which may imply conserved structural or functional characteristics between R1 and R5 pyocins. Based on this, we tested the inhibitory effects of R1 pyocins (using filtrates from ST281/M0027 and ST260/M0089) on our clinical isolates (Table S1). Surprisingly, most ST111 strains were not inhibited by R1 pyocin. We also tested the susceptibility of PA14 LPS mutants to filtrates from R1 (ST260/M0089), R2 (ST549/PAO1), and R5 (ST111/218M0087) pyocin-encoding strains. CFU results and growth kinetics confirmed that PA14*ΔwaaL*, PA14*wbpL* and PA14*wbpM* were sensitive to R1 and R5 filtrates, but resistant to R2 filtrate (Fig 5C and Figs S8B and S8C).

We considered two potential explanations for the different effects seen for R1, R2 and R5 pyocins. First, ST253/PA14 may possess a detoxification system that provides defense against the R2 pyocin it produces, but that is ineffective against other R pyocins. Second, WaaL activity may specifically preclude the binding of R1 and R5 pyocins. To test these hypotheses, we first selected a few clinical isolates that encode R1 (ST260/M0089, ST179/MB3221), R2 (ST447/MB3216, ST856/MB3340), or R5 (ST111/218M0087, ST111/M0101, or ST446/M0186) pyocins, along with two lab-adapted R2 pyocin-encoding strains (ST253/PA14 or ST549/PAO1). Production of R pyocins was confirmed using a CFU assay with the R pyocin indicator strain 13s (Fig 6A).

**Figure 6.**
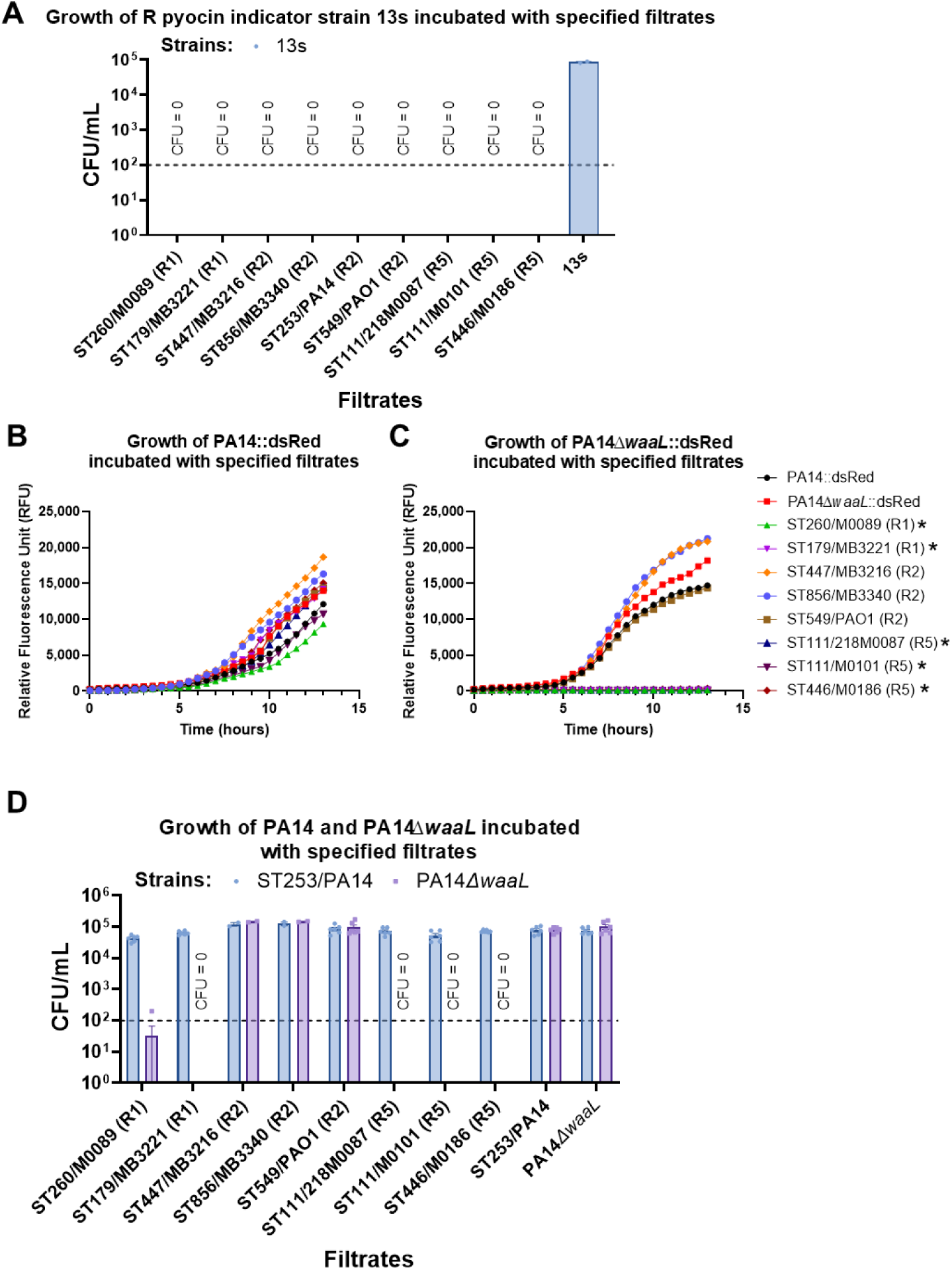
PA14*ΔwaaL* is sensitive to R1 and R5 pyocins. (**A**) CFUs (colony-forming units) of R pyocin indicator strain 13s after 1 hour incubation with filtrates from self, R1 pyocin-encoding strains ST260/M0089, ST179/MB3221, R2 pyocin-encoding strains ST447/MB3216, ST856/MB3340, ST253/PA14, ST549/PAO1, and R5 pyocin-encoding strains ST111/218M0087, ST111/M0101, or ST446/M0186. (**B-C**) Growth of the PA14::dsRed (**B**) and PA14*ΔwaaL*::dsRed (**C**) incubated with filtrates from self, R1 pyocin-producing strains ST260/M0089, ST179/MB3221, R2 pyocin-producing strains ST447/MB3216, ST856/MB3340, ST549/PAO1, and R5 pyocin-producing strains ST111/218M0087, ST111/M0101, or ST446/M0186. Three biological replicates were performed. Filtrates labeled with asterisk (*) exhibited strong growth inhibition of PA14*ΔwaaL*::dsRed. (**D**) CFUs (colony-forming units) of PA14 and PA14*ΔwaaL* after 1 hour incubation with filtrates from self, R1 pyocin-producing strains ST260/M0089, ST179/MB3221, R2 pyocin-producing strains ST447/MB3216, ST856/MB3340, ST549/PAO1, and R5 pyocin-producing strains ST111/218M0087, ST111/M0101, or ST446/M0186. For (**A,D**), detection limit of CFU assay (horizontal dashed line) is 100 colony/mL per individual technical replicate.

Next, we challenged PA14 and PA14*ΔwaaL* with filtrates from the selected R1-, R2-group-, and R5-producing strains (Figs 6B-D). To decrease bias in our panel, R1 and R2 pyocin-encoding strains from a separate site (MD Anderson Cancer Center, Houston, Texas) were added (strains with MB designation). Growth of a dsRed-engineered PA14*ΔwaaL* mutant was significantly inhibited by all filtrates from R1 and R5 pyocin-producing strains, compared with that of a wild-type, dsRed-expressing PA14 (Fig 6B and 6C, labeled with an asterisk). Further quantification via CFU assay confirmed the strong growth inhibition exhibited by these filtrates (Fig 6D). These data support the hypothesis that deficiency in LPS biosynthesis is associated with sensitivity towards R1 and R5 pyocins. To test whether the production of R2 pyocin in PA14 contributes to the resistance of PA14*ΔwaaL* to R2 pyocin, we then deleted *waaL* in the PA14*Δpyocins* background and challenged the resulting strain, PA14*ΔpyocinsΔwaaL*, with the same panel of filtrates. Interestingly, the mutant remained resistant to R1 filtrates, but became sensitive to R2 and R5 filtrates (Fig 7A; see Fig S9A for the second, independently-derived, line of PA14*ΔpyocinsΔwaaL*). This outcome may be partially explained by the deletion of 31 genes for both R and F pyocins (42) (see Discussion).

**Figure 7.**
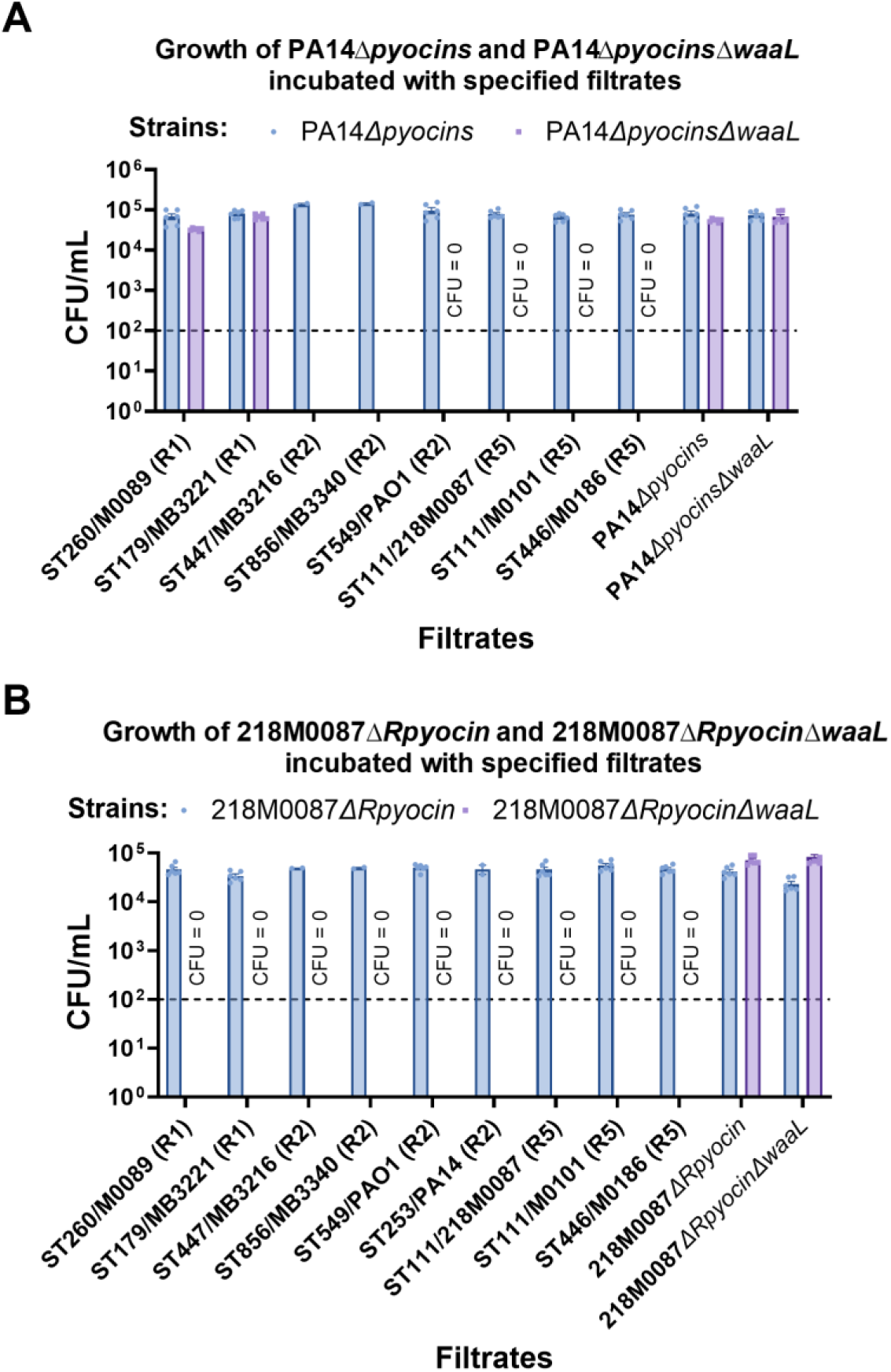
*waaL* deletion expands the susceptibility range of *P. aeruginosa* to R pyocins. (**A**) CFUs (colony-forming units) of PA14*Δpyocins* and PA14*ΔpyocinsΔwaaL* after 1 hour incubation with filtrates from self, R1 pyocin-producing strains ST260/M0089, ST179/MB3221, R2 pyocin-producing strains ST447/MB3216, ST856/MB3340, ST549/PAO1, and R5 pyocin-producing strains ST111/218M0087, ST111/M0101, or ST446/M0186. (**B**) CFUs (colony-forming units) of 218M0087*ΔRpyocin* and 218M0087*ΔRpyocinΔwaaL* after 1 hour incubation with filtrates from self, R1 pyocin-producing strains ST260/M0089, ST179/MB3221, R2 pyocin-producing strains ST447/MB3216, ST856/MB3340, ST253/PA14, ST549/PAO1, and R5 pyocin-producing strains ST111/218M0087, ST111/M0101, or ST446/M0186. For (**A-B**), detection limit of CFU assay (horizontal dashed line) is 100 colony/mL per individual technical replicate.

To further investigate the impact of the *waaL* deletion in non-R2 strains, we constructed the following deletions in an R5-expressing strain: 218M0087*ΔwaaL* and 218M0087*ΔRpyocinΔwaaL*. All lines of 218M0087*ΔwaaL* tested displayed strong growth deficiency, likely due to sensitivity to its own R5 pyocin and hence were excluded from further experimentation. 218M0087*ΔRpyocinΔwaaL* bacteria, which did not produce R5 pyocin, were challenged with the same panel of filtrates. Surprisingly, both lines of this mutant were sensitive to all filtrates containing R pyocins (Fig 7B and Fig S9B). Overall, the *waaL* deletion caused susceptibility of previously-resistant strains to R5 pyocins, but sensitivity of these mutants to R1 or R2 pyocins was context-dependent.

### Most global high-risk *P. aeruginosa* clones encode R5 pyocin

To more broadly examine the relationship between R pyocin subtype and epidemiological risk associated with a given ST type, we analyzed data from the Pseudomonas Genome Database (PGD) which includes 5,135 *P. aeruginosa* strains (Fig 8A) (49). These strains were isolated from myriad global sources, including human and animal infections, hospital facilities, and natural environments, providing a relatively comprehensive representation. 33.6% (1,726/5,135) of isolates harbored genes for R5-type pyocin tail fiber, which encode a distinct peptide sequence compared to R1 or R2 proteins (Fig S10A). A phylogenetic tree based on the tail fiber protein sequences from the isolates used in this study revealed the existence of several subclades (Figure S10B).

**Figure 8.**
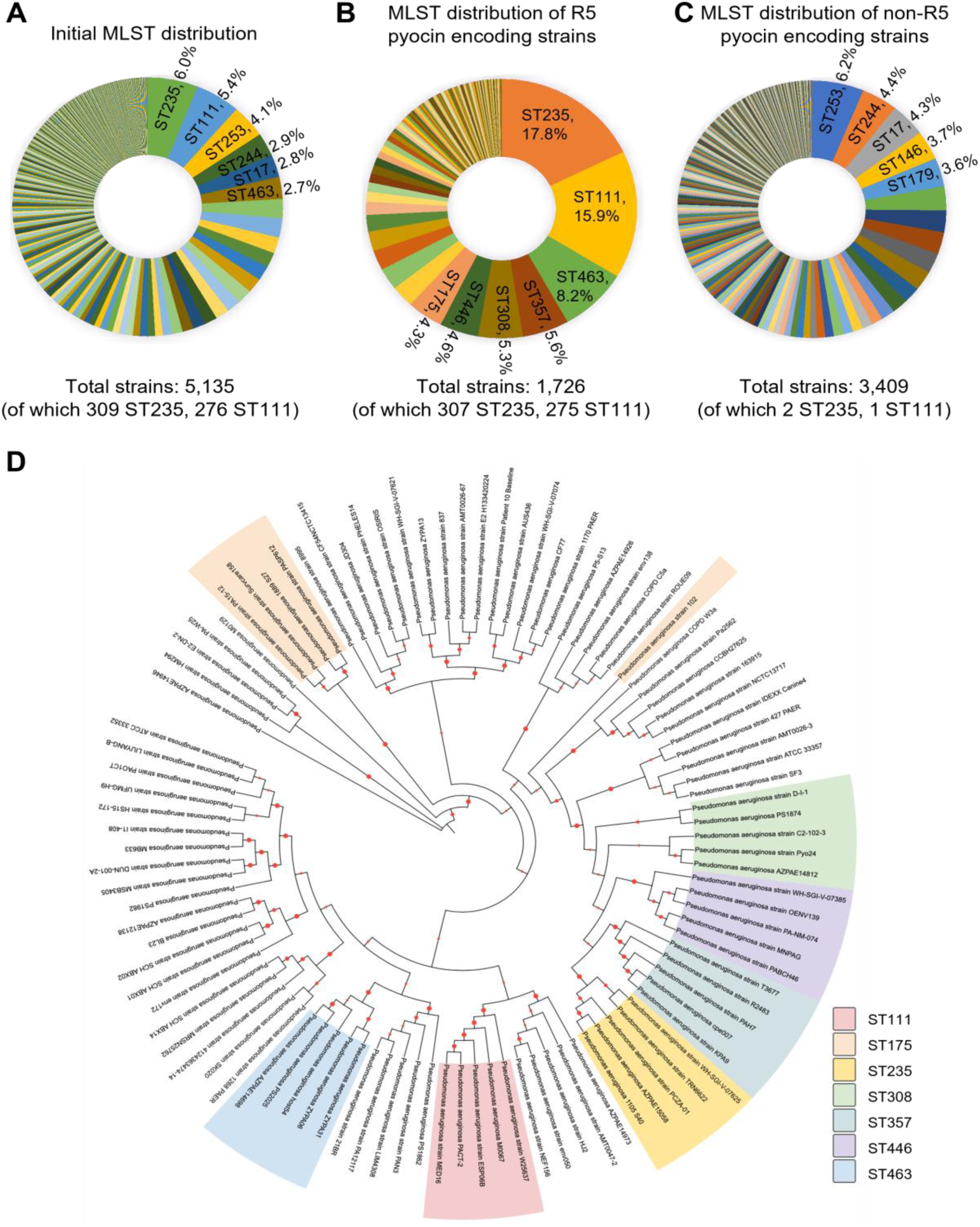
International high-risk sequence types are enriched in R5 pyocin coding strains. (A-C) Pie chart of input MLST distribution (**A**), MLST distribution of R5 pyocin carriers (**B**), MLST distribution of non-R5 pyocin carriers (**C**). (**D**) A phylogenetic tree of 100 isolates from the 20 most abundant MLSTs in 5,135 isolate set.

Multiple globally-prevalent, high-risk sequence types encoded the R5 pyocin tail fiber gene. ST235, the most prevalent high-risk sequence type (50, 51) comprised 17.8% of the R5 pyocin-encoding strains. ST111 (17) was 15.9%, ST463 (52) was 8.2%, ST357 (19) was 5.6%, ST446 (53) 5.3%, ST308 (54) was 4.6%, and ST175 (55) was 4.3% of isolates with the R5 pyocin (Fig 8B). Although the majority of high-risk sequence types encoded R5 pyocin genes (56), this association was not universal. For example, most strains in the ST244 group (the fourth most common sequence type in the PGD) did not encode R5 pyocins (Fig 8A and C). ST244 is also an epidemic, high-risk clone of *P. aeruginosa* (19), suggesting that R pyocin subtype is not the sole determinant of strain dominance.

To examine whether R5 pyocin production is enriched in a cluster of closely-related sequence types, we analyzed the phylogenetic relationship of the 20 most abundant MLSTs within the 5,135 isolates in the PGD, including 7 MLSTs of R5 pyocin carriers and 13 MLSTs of non-R5 pyocin carriers. Five isolates were randomly selected from each R pyocin sequence type (100 isolates total). We constructed a phylogenetic tree using 100 randomly-selected genes using the Codon Tree method (57). Analysis showed that strains belonging to the same MLST tended to cluster together (Fig 8D). A clade of isolates carrying an R5 pyocin-encoding gene (including isolates from ST235, ST357, ST446, and ST308) was observed, implying greater relation between the strains than merely the presence of the R5 pyocin gene. These R5 coding strains diverged from other strains more recently, suggesting that the capacity for R5 pyocin production could confer an evolutionary advantage.

However, other R5 pyocin-encoding sequence types, including ST111, ST463, or ST175, did not appear to be closely related to the clade of R5 pyocin-encoding MLSTs described above or to each other. This may indicate that they acquired R5 genes independently, perhaps via horizontal gene transfer.

In order to establish the non-random nature of the association between R5 pyocin and internationally-recognized, high-risk strains, we conducted an in-depth analysis of MLST distributions amongst R1-4 pyocin coding strains. Due to high sequence similarity amongst R2, R3, and R4 pyocins (over 98.84%), the MLST analysis treated R2, R3, and R4 as one category. Results showed that none of the R1 or R2-4 pyocin coding strains were found to belong to ST235 or ST111 (Fig S11). Together, these collective findings strongly suggest that R5 pyocin production could indicate the likelihood of a newly emerging strain of *P. aeruginosa* to be a high-risk clone.

## Discussion

Previously, we showed that carbapenem non-susceptible ST111 *P. aeruginosa* clinical isolates from HM/HCT patients at OHSU exhibited a fitness advantage over the laboratory reference strain ST253/PA14, in part due to OprD deficiency (17). However, OprD deficiency failed to explain ST111’s dominance over other OprD-deficient clinical isolates of *P. aeruginosa* from the same patient population. In this study, we observed strong inhibition of growth of certain non-ST111 strains, including ST260/M0089, ST291/M0103, ST233/M0104, and ST299/M0128, by ST111 and ST253/PA14. This phenomenon depended upon a high-molecular weight, heat-labile, material produced by ST111 or ST253/PA14. High-throughput screening of transposon mutants identified this factor as an R-type pyocin, and further analysis indicated that all of the ST111 isolates encoded R5 subgroup pyocins (unlike PA14 and PAO1, which encode R2 pyocin). R5 subgroup pyocins display the broadest bactericidal activity against competing bacteria. This work represents, to the best of our knowledge, the first to demonstrate an association between R-pyocin and strain dominance in a clinical setting.

Without functional WaaL, LPS is bereft of O-antigen, which leaves the R pyocin receptor in the LPS core accessible (58). Interestingly, deletion of *waaL* in the ST253/PA14 background sensitized the mutant to R1 and R5 pyocins, but not its own or ST549/PAO1’s R2 pyocin. This may be due to the presence of the tail fiber assembly protein in PA14 R2-subtype pyocin operon. Production of excess tail fiber assembly protein, which is predicted to directly bind to R2-subtype tail fiber, could limit the potential for bacterial cytotoxicity. Assembly proteins have divergent protein sequences, which may prevent the R2 assembly protein from interacting with R1 or R5 pyocins, but would likely protect against R2-4 pyocins, which have strong sequence similarity. Deletion of operons for both R and F pyocins, along with deletion of *waaL*, rendered PA14*ΔpyocinsΔwaaL* susceptible to R2 and R5 pyocins, while deletion of *waaL* in the 218M0087*ΔRpyocin* mutant sensitized it to all R pyocin-containing filtrates. This suggests that deficiency of the O-antigen ligase WaaL most commonly rendered *P. aeruginosa* susceptible to R5 pyocins. Other proteins may be involved in determining the sensitivity of *P. aeruginosa* LPS mutants to R pyocins from other clades. Importantly, some of the R5 pyocin-sensitive strains lacked obvious mutations in *waaL, wbpL*, or *wbpM*, suggesting that other factors may also influence pyocin sensitivity.

All known binding sites for R pyocins are on the LPS outer core, but there are differences in reported binding sites (26). R1 pyocin binds the first L-rhamnose and R2 pyocin binds the first α-glucose residue in LPS, while R5 binds the second α-glucose residue (26). This difference may drive the broader killing spectrum for R5 pyocins, but additional research is necessary. In addition to serving as the target for R pyocins, LPS is a pathogen-associated molecular pattern recognized by the human innate immune system (59). Future work may investigate whether there is a relationship between changes in LPS that facilitate immune avoidance and those that alter susceptibility to pyocins.

Finally, previous explanations for the dominance of *P. aeruginosa* MLSTs like ST111 have focused on their propensity for acquiring multi-drug resistance in clinical settings. This explanation seems insufficient when dominant and minority sequence types have similar or identical resistance profiles. Our research suggested that pyocin subtype may be an additional determinant of clinical strain dominance. 5,135 strain-typed *P. aeruginosa* isolates collected world-wide were examined for their pyocin subtype. Notably, several high-risk STs, including ST235 and ST175, which together with ST111 are regarded as the three major international high-risk sequence types (20), contain the genes that are related to R5 production. Importantly, these three sequence types are not closely phylogenetically-related (see Fig 8). Our results suggest that some high-risk sequence types may reduce the viability of competing strains by producing R5 pyocins, which have the broadest killing spectrum. The association between high-risk sequence types and R5 pyocins may justify considering R5 pyocin production amongst the risk determinants for *P. aeruginosa* strains and supports calls for typing *P. aeruginosa* isolates by pyocins (60). Our phylogenetic analysis also underscores the prominent role of R5 pyocin-encoding strains in the *P. aeruginosa* community, showcasing their evolutionary significance. This insight helps elucidate their global prevalence and the associated public health risks. While R5 pyocin may not be the sole determinant for predominance, our data strongly suggest it confers a substantial competitive advantage.

## Materials and methods

### Bacterial strains and growth conditions

*Escherichia coli* SM10 or OP50, or *Pseudomonas aeruginosa* were grown in Luria-Bertani Lennox broth (LB, 10 g/L tryptone, 5 g/L yeast extract, 5 g/L NaCl) or on LB agar plates fortified with 1.5% agar at 37°C. *Enterococcus faecalis* OG1RF was grown in Brain Heart Infusion broth (BHI). When appropriate, 25 μg/mL irgasan (to specifically select for *P. aeruginosa*); or 30 or 15 μg/mL gentamicin; or 1.5 µg/mL imipenem; or 50 µg/mL rifampicin; or 8 or 16 µg/mL meropenem; or 300 µg/mL carbenicillin was added to liquid or solid media. Markerless deletions were generated using the pEXG2 vector with counterselection on no-salt LB plates containing 20% sucrose (10 g/L tryptone, 5 g/L yeast extract, 20% (w/v) sucrose, 15 g/L agar). Transposon-insertion mutants came from commercially-available libraries (38, 61). The *P. aeruginosa* strains used in this study are summarized in Table 5.

**Table 5.** *Pseudomonas aeruginosa* strains used in this study.

| Strain | MLST | Genotype or description | Source or reference |
| --- | --- | --- | --- |
| PA14 | 253 | Laboratory wild-type strain of *P. aeruginosa* | Laboratory stock (62) |
| PAO1 | 549 | Laboratory wild-type strain of *P. aeruginosa* | Laboratory stock (62) |
| M0067 | 111 | *P. aeruginosa* bloodstream clinical isolates obtained from the OHSU Clinical Microbiology lab | (17) |
| M0101 | 111 |  |  |
| 218M0087 | 111 |  |  |
| M0025 | 111 |  |  |
| M0134 | 111 |  |  |
| M0169 | 111 |  |  |
| M0177 | 111 |  |  |
| M0249 | 111 |  |  |
| M0003 | 111 |  |  |
| M0117 | 446 |  |  |
| M0186 | 446 |  |  |
| M0122 | 308 |  |  |
| M0162 | 281 |  |  |
| 075M0106 | 384 |  |  |
| M0068 | 132 |  |  |
| M0103 | 291 |  |  |
| M0104 | 233 |  |  |
| M0128 | 299 |  |  |
| M0013 | 17 |  |  |
| M0027 | 281 |  |  |
| M0043 | 308 |  |  |
| M0089 | 260 |  |  |
| MB1854 | 244 | *P. aeruginosa* bloodstream clinical isolates obtained from the clinical microbiology laboratory at MD Anderson Cancer Center, Houston, TX. | This study |
| MB3221 | 179 |  |  |
| MB3090 | 298 |  |  |
| MB2853 | 298 |  |  |
| MB3613 | 235 |  |  |
| MB3244 | 111 |  |  |
| MTC2191 |  | 13s (R pyocin indicator) | Originally published in (41), and gifted by Cabeen Lab at Oklahoma State University (42) |
| MTC2192 |  | PML1516d (S pyocin indicator) | Originally published in (63), and gifted by Cabeen Lab at Oklahoma State University (42) |
| MTC2326 | 253 | PA14Δ*pyocins* (Δ07970-08300) | Gifted by Cabeen Lab at Oklahoma State University (42) |
| | 253 | PA14Δ*pyocins* Δ*waaL* | This study |
| | 253 | PA14Δ*waaL* | This study |
| | 253 | PA14Δ*flgK* | Gifted by O'Toole Lab at University of Dartmouth (47) |
|  | 291 | M0103::*Carb<sup>R</sup>* (M0103::*dsRed*) | This study |
|  | 233 | M0104::*Carb<sup>R</sup>* (M0104::*dsRed*) | This study |
|  | 299 | M0128::*Carb<sup>R</sup>* (M0128::*dsRed*) | This study |
|  | 253 | PA14::*dsRed* | (17) |
| | 253 | PA14Δ*waaL*::*dsRed* | This study |
| | 111 | 218M0087Δ*Rpyocin*(ΔPA14_08090 ortholog gene) | This study |
|  | 253 | PA14*waaL* | (38) |
|  | 253 | PA14*galU* | (38) |
|  | 253 | PA14*gmd* | (38) |
|  | 253 | PA14*lptD* | (38) |
|  | 253 | PA14*lpxC* | (38) |
|  | 253 | PA14*lpxO1* | (38) |
|  | 253 | PA14*lpxO2* | (38) |
|  | 253 | PA14_69170 | (38) |
|  | 253 | PA14*pagL* | (38) |
|  | 253 | PA14*wzm* | (38) |
|  | 253 | PA14*wzt* | (38) |
|  | 253 | PA14*wzz* | (38) |
|  | 253 | PA14*wbpM* | (38) |
|  | 253 | PA14*wbpZ* | (38) |
|  | 253 | PA14*wbpL* | (38) |
|  | 253 | PA14*wbpY* | (38) |
|  | 253 | PA14*wbpW* | (38) |
|  | 253 | PA14*wbpX* | (38) |

529 <sup>OE</sup>: overexpression; Tn: transposon insertion

### Caenorhabditis elegans strains

A temperature-sensitive sterile strain of *C. elegans glp-4*(*bn2*) was used in this project (64). Worms were maintained on nematode growth media (NGM) plates seeded with *E. coli* OP50. For propagation, *glp-4*(*bn2*) worms were incubated at 15 ℃. For experiments, synchronized L1 larvae of *glp-4*(*bn2*) worms were grown overnight on concentrated OP50 lawn at room temperature, and then shifted to 25°C to induce sterility for 44∼48 h prior to use.

### *In vitro* or *in vivo* competition assays

These assays were performed as previously described (17). In brief, for *in vitro* assay, two *P. aeruginosa* were cultured overnight separately in LB medium. The next day, *P. aeruginosa* was pelleted and washed with 1 mL sterile water in a 1.5 mL Eppendorf tube to remove secreted materials, then resuspended in 1 mL sterile water. Two *P. aeruginosa* isolates were mixed 1:1 based on the OD600, or with an equivalent volume of sterile water instead for single-strain control groups, and spread onto SK plates. 0 h or after 24 h of incubation at 37°C, bacteria were diluted 1000x before additional serial 10-fold dilutions in a 96-well plate. Serial dilutions were then seeded on the appropriate antibiotic plates: meropenem (8 μg/mL, Combi-Blocks) for carbapenem-resistant clinical isolates or rifampicin (50 μg/mL, Fisher Scientific) for ST253/PA14, or gentamicin (15 μg/mL, Fisher Scientific) for ST253/PA14 transposon-insertion mutants. If two clinical isolates with carbapenem resistance were examined, seeding on a plain LB plate yielded total colonies while seeding on 300 μg/mL carbenicillin (Fisher Scientific) selected for the strain transformed with the plasmid with carbenicillin resistance. The colonies of the second strain were then calculated by subtracting colonies on the carbenicillin LB plate from those on the no-antibiotic LB plate. After incubation overnight, colonies were counted. Three biological replicates were performed with three technical replicates each.

For *in vivo* assay, two *P. aeruginosa* were mixed 1:1 based on the OD600 and spread onto SK plates as mentioned in *in vitro* competition. The initial ratio of two *P. aeruginosa* was determined at this time point by serial dilutions and appropriate antibiotic selection. The mixed bacteria were incubated 24 h at 37 °C, then another 24 h at 25 °C. Young adult *glp-4*(*bn2*) worms were picked onto the bacterial lawn and left for 40 h of infection at 25 °C. 15 adult worms were then picked into a 1.5 mL tube with S Basal buffer containing levamisole to prevent pumping, washed six times, and lysed via vortexing with zirconium beads (Fisher Scientific, 1.0 mm). Buffer from the final wash (blank control) and worm lysate were serially diluted 5-fold and seeded onto the appropriate antibiotic plate. Three biological replicates were performed with three technical replicates each.

### Growth kinetics

A single *P. aeruginosa* colony was inoculated into 5 mL of LB liquid broth and was incubated at 37 ℃ for 12∼16 h. The next day, a final OD600 0.1 was achieved by diluting overnight cultures in LB. A 96-well plate containing 100 μL final solution in each well was covered with an air-permeable membrane and placed into Cytation5 multimode plate reader (BioTek) for running growth kinetics at 37 ℃. OD600 measurements were obtained every 30 minutes. Three biological replicates were performed with three technical replicates each.

### Filtrate assay

A single colony was inoculated into 5 mL of LB liquid broth and was incubated at 37 ℃ for 12∼16 h. The overnight culture was spun down at 10,000 g for 1 minute at room temperature. The supernatant was harvested and passed through a 0.22 μm syringe filter to remove any remaining cells, yielding bacteria-free spent media (filtrate). Heated filtrate was obtained by incubating 1 mL of filtrate at 95℃ for 30 minutes in a dry bath. A 100 kDa centrifugal filter (Merck Millipore) was used to separate small and large molecular fractions of the filtrate. A total of 40 mL of filtrate was loaded into the top part of the centricon and was centrifuged at 3,000 g for 30 minutes. The filter was washed with 40 mL of PBS. 200 μL filtrate harvested from the centricon was diluted in 2 mL PBS. The growth kinetics were performed as previously described with a final OD600 0.1 in 50% filtrate. Three biological replicates were performed with three technical replicates each.

### Bactericidal effect test

For CFU assay, a single *P. aeruginosa* colony was inoculated into 5 mL of LB liquid broth and was incubated at 37 ℃ for 14 h. The overnight culture was diluted to ∼ 10^5^ colony/mL, then 1:1 mixed with prepared filtrate (100 μL in total) in 96-well plate and incubated at 37°C. After 1 h of incubation with filtrate, a 10 μL of well content was sampled. This aliquot was serially diluted 5-fold and seeded to a LB plate. Colonies were counted the following day and CFU/mL was calculated. Three biological replicates were performed each.

For imaging, combined ST233/M0104 diluted culture and filtrate were prepared as previously described. Filtrate and diluted culture were combined 1:1 for 1 mL total volume and incubated with shaking at 37°C. After incubation for three hours at 37 °C, the culture was stained with 40 μM acridine orange and 1 μg/mL propidium iodide, then incubated for another hour. The culture was then pelleted by centrifugation at 10,000 g for 10 minutes, and the pellet was resuspended in S Basal before dropping on an agar-padded microscope slide. Images were taken using a Zeiss ApoTomeM2 Imager fluorescent microscope (Carl Zeiss, Germany) with a 40x oil objective magnification. Three biological replicates were performed.

### PA14 transposon insertion mutant library screen

For the primary screen, library plates were stamped into a 96-well plate with 100 μL LB and incubated overnight at 37 °C to create sub-cultures. Sub-cultures were then used to inoculate a 96-deep-well plate with fresh 1 mL LB for a second overnight incubation in a Multitron Pro shaking incubator (Infors HT) at 37 °C. 50 μL of the deep well cultures were incubated in a 96-well plate with meropenem (16 μg/mL, Combi-Blocks) for 5 hours, yielding a “suppressed culture.” An overnight culture of M0104::dsRed was diluted to an OD600 of 0.1 and combined 1:1 by volume with the suppressed cultures in a 96-well plate. The 96-well plates were covered with a lid and incubated at 37°C. After 0, 6, or 18 h, plates were placed into a Cytation5 multimode plate reader to measure OD600 and dsRed fluorescence. Two technical replicates were performed per library plate. Hits were defined as having fluorescence above the plate average plus 3 standard deviations at 6 and/or 18 hours in both replicates. Wells were excluded if 0-hour OD600 indicated an absence of PA14 library mutant growth. For the secondary screen, hits were tested using the filtrate assay as described previously. Three biological replicates were performed with two technical replicates each.

### Pyocin indicator assays

The filtrate was obtained as previously described. Indicator strains were grown in 5 mL of LB liquid at 37℃ for 12∼16 h. OD600 of the overnight culture OD600 was measured. The growth kinetics experiments were performed as previously described with a final OD600 0.1 in 50% filtrate. Three biological replicates were performed with three technical replicates each.

### *P. aeruginosa* gene deletion generation

Two 500-600 bp homology regions flanking the gene of interest were cloned via PCR and were inserted into the *Xba*I/*Sac*I pretreated pEXG2 vector via Gibson assembly (New England Biolabs). 2 μL of the product were transformed into 100 μL *E. coli* DH5**α** competent cells via heat shock method (42°C for 40 seconds, cool down on ice for 2 minutes). Cells were recovered in 1 mL SOC medium by incubation in a shaker (200 rpm) for 2 hours at 37 ℃ and then were spread onto 12.5 μg/mL gentamicin selective LB agar plates for colony growth. Colonies were tested via colony PCR. Plasmids from selected colonies were sequence-verified and transferred into *E. coli* SM10 competent cells via heat shock method. The SM10 cells containing knockout plasmid were grown overnight in 5 mL LB broth with 12.5 μg/mL gentamicin added. The *P. aeruginosa* strain of interest was incubated overnight in 5mL LB broth without gentamicin. The cell pellets of SM10 and the *P. aeruginosa* strain of interest were harvested via centrifugation at 10,000 g for 1 minute, mixed in 50 μL of LB broth, and spotted onto LB agar plate for conjugation, 6 hours at 37 °C. After incubation, the mixed cells were washed from the plate with 1 mL LB. 20-50 μL of bacteria were added to 1 mL fresh LB and spread onto an irgasan (25 μg/mL in ethanol)/gentamicin (12.5 μg/mL in water) LB plate, incubating at 37°C overnight. That allowed for selections of *P. aeruginosa* colonies with the recombinant plasmid. To remove the plasmid from *P. aeruginosa*, a colony from the irgasan/gentamicin plate was inoculated into a 5 mL LB medium without antibiotics. The overnight culture was then streaked onto a counter-selection plate (10 g/L tryptone, 5 g/L yeast extract, 20% (w/v) sucrose, 15 g/L agar) and incubated overnight at 37°C. Next day, ten colonies were selected to check the gene deletion via the single colony PCR. At least two independent lines were isolated. At least one of them was sequence verified.

### Whole genome sequence analysis of constructed deletion mutants

Bacterial genomic DNA of PA14 was purified from overnight culture using FastPure DNA Isolation Mini Kit (Vazyme). Paired end Illumina whole genome sequencing was performed by the SEQCENTER for 400 minimum read counts per sample. To verify the deletion, raw sequencing reads from each mutant were compared with the respective reference genome using breseq (65, 66).

### Phage plaque assay

The receptor (ST291/M0104 in this study) and the effector (ST253/PA14, ST111/218M0087, and ST111/218M0087*ΔRpyocin* in this study) strains were grown in 5 mL of LB liquid broth at 37°C until the stationary phase was reached (about 12-16 hours). Next day, 200 μL of the receptor was added into 4 mL of fresh LB broth, incubating in a shaker (200 rpm) at 37°C for 3 hours of growth to the log phase. The effector filtrate was prepared as described in the filtrate assay. To make 0.5% agar LB medium, the mixture of 0.25 g agar and 50 mL LB was heated in a microwave oven until all agar dissolved and then cooled down in a 45°C water bath. 5 mL of 0.5% agar LB, 100 μL of the log-phase receptor, and 10 μL or 200 μL of the filtrate from the effector mixed well and poured onto 1.5% LB agar plate with swirling the plate around to properly dispense the contents. The receptor (ST291/M0104 in this study) filtrate was used as a blank control. After solidifying, plates were incubated at 37°C overnight. The plaques on the plates were checked and imaged the following day.

### Swimming motility assay

Swimming motility assay was performed as previously described (47). Briefly, a single *P. aeruginosa* colony was inoculated into 5 mL of LB liquid broth and was grown at 37 ℃ for > 10 h. Swimming agar was prepared on the day of the assay by supplementing 0.3% M8 agar with 0.2 % glucose, 0.5 % casamino acids, and 1 mM MgSO4. A sterile disposable pipette tip was used to dip into overnight culture and then stab halfway into the agar layer of the plate. The agar plates were incubated upright at 37 °C for 16 h to observe the phenotype. Swimming motility was measured by the diameter of bacterial radial growth.

### RNA extraction

A 50 μL pellet of *P. aeruginosa* cells was collected from the overnight culture and resuspended in 150 μL sterile water in a 1.5 mL microcentrifuge tube. 600 μL of Trizol was mixed with *P. aeruginosa* by pipetting up and down several times. The tube was stored at -80°C for at least 5 hours and then was taken out from the freezer and vortexed until the iced pellet melted. With another 400 μL of Trizol added, cells were vortexed for 10 seconds and set on the ice for 5 minutes. The procedure was repeated after 150 μL of BCP (1-Bromo-3-chloropropane) was added. The cells’ debris was spun down at the maximum speed at 4°C for 20 minutes. The supernatant was transferred into a new prechilled 1.5 mL tube. 1:1 v/v isopropanol was added to the supernatant. After 5 minutes of incubation at room temperature, the RNA was spun down at 12,000 rpm for 10 minutes at 4°C. The RNA pellet was washed with 1 mL prechilled 80% ethanol twice and then dissolved in DEPC water. This RNA was used for quantitative RT-PCR. Three biological replicates were performed.

### Quantitative RT-PCR

cDNA was synthesized using a cDNA synthesis kit (New England BioLabs). Quantitative reverse-transcription real-time PCR (qRT-PCR) was conducted in a CFX-96 real-time thermocycler (Bio-Rad) using SYBR green AzuraQuant Fast Green Fastmix(Azura). Fold changes were calculated using a △Ct method using *gyrB* (DNA gyrase subunit B) as a housekeeping gene. The primers used in qRT-PCR are shown in Table 6.

**Table 6.**
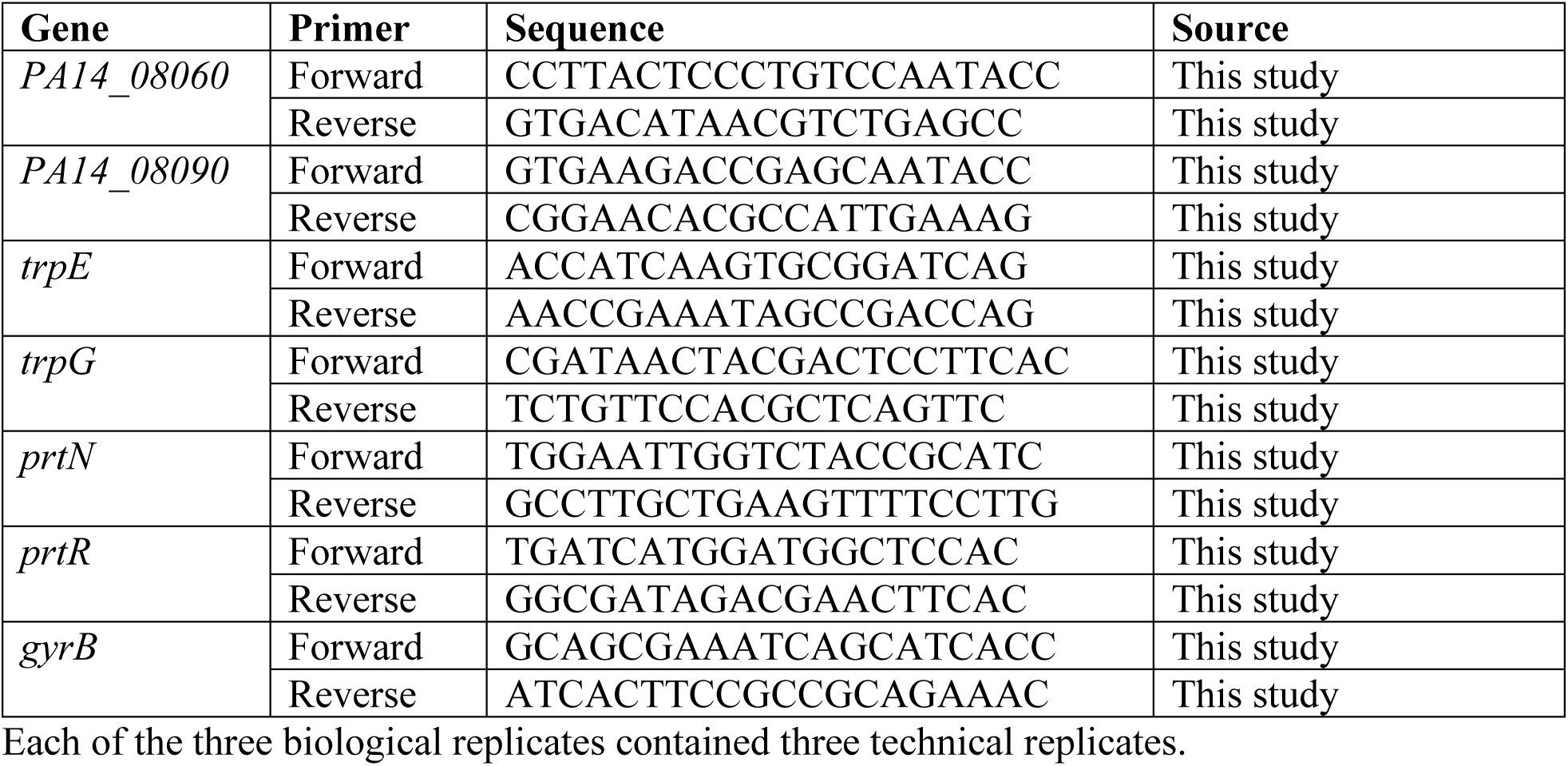
Quantitative PCR primers.

### MLST distribution analysis

The amino acid sequence and MLST information of 5,135 *P. aeruginosa* isolates were collected from the Pseudomonas Genome Database (https://www.pseudomonas.com). Specifically, the amino acid sequences of 7,465 *P. aeruginosa* isolates were downloaded and strain information, including strain name, host, isolation source, location, and MLST, was downloaded from the website. Strains that did not have MLST information or the strains whose names couldn’t match to any file name of the protein sequence were removed, resulting in a total of 5,135 isolates left for pyocin analysis. Several unique fragments were used to distinguish the subtype of R pyocin: two fragments (DHPGGIIDR, VSVSNTGCVIVSSEYYGLAQNYG) for R1 pyocin, two fragments (TCPADADASI, FRGATTTTAVIRNGYFAQAVLSWE) for R2&3&4 pyocin, and five fragments (SNPSTLA, AVSTTSDP, PSARDWLPWKRC, GGSFSK, GKPATFPPT) for R5 pyocin.

### Phylogenetic tree construction

A total of 100 isolates, which were randomly selected from the 20 most abundant MLSTs in 5,135 isolates with 5 strains for each MLST, were employed to perform the phylogenetic analysis. The genomic sequence of 100 isolates was used to construct the phylogenetic tree on the website of Bacterial and viral bioinformatics resource center (BV-BRC: https://www.bv-brc.org/app/PhylogeneticTree) (67). As the default setting, 100 genes were compared to yield the phylogenetic tree. The .nwk file with strain names was downloaded from the BV-BRC and was further annotated on the website iTOL (https://itol.embl.de).

## Supporting information

Combined Sup Materials

## Supporting information

**Figure S1. Filtrate from ST111 and ST446 strains impaired the growth of non-ST111 strains.** (**A-C**) Growth of non-ST111 strains (ST233/M0104, ST291/M0103, ST299/M0128) incubated with 50% filtrates from self or ST111 strains (ST111/M0067, ST111/M0101, ST111/218M0087). (**D-F**) Growth of non-ST111 strains (ST233/M0104, ST291/M0103, ST299/M0128) incubated with 50% filtrates from self or ST446 strains (ST446/M0117, ST446/M0186), respectively. Three biological replicates were performed with three technical replicates for each.

**Figure S2. A large, heat-sensitive complex exhibits rapid bactericidal activity toward non-ST111 strains.** (**A-C**) Growth of non-ST111 strains (ST233/M0104, ST291/M0103, ST299/M0128) incubated with 50% filtrates from self or untreated, heated, or fractionated ST111/M0101. (**D-F**) Growth of non-ST111 strains (ST233/M0104, ST291/M0103, ST299/M0128) incubated with 50% filtrates from self or untreated, heated, or fractionated ST253/PA14 filtrates. (**G**) Images of ST233/M0104 cells stained with acridine orange (AO, all cells) and propidium iodide (PI, dead cells) after 4 hours of incubation with self *vs.* ST253/PA14 filtrate. Three biological replicates were performed with three technical replicates for each. Scale bar: 25 μm.

**Figure S3. ST233/M0104::dsRed growth curve in mock library screen reproduces filtrate results.** (**A, B**) ST233/M0104::dsRed growth curves, OD600 (**A**) and red fluorescence-based (**B**), in the presence of 50% filtrate from ST233/M0104 (self-control), ST253/PA14, *E. coli* OP50, or *E. faecalis* OG1RF. (**C, D**) ST233/M0104::dsRed growth curves, OD600 (**C**) and red fluorescence-based (**D**), in the presence of ST253/PA14, *E. coli* OP50, or *E. faecalis* OG1RF, each with or without 8 μg/mL meropenem. Two biological replicates were performed with four technical replicates for each using primary screen protocol.

**Figure S4. Screen hits display decreased growth inhibition of the sensor strains.** (**A-C**) Growth of ST233/M0104::dsRed (**A**), ST291/M0103::dsRed (**B**), or ST299/M0128::dsRed (**C**) in the presence of filtrates from self (negative control), WT ST253/PA14 (positive control), or three strong PA14 Tn mutant hits in the secondary screen. Three biological replicates were performed with two technical replicates for each.

**Figure S5. Screen hits do not inhibit the growth of R pyocin indicator strain.** (**A**) Growth of 9 confirmed strong screen hits compared to WT PA14 (control). (**B**) Growth of R pyocin indicator strain 13s with filtrates from self, ST253/PA14, PA14*Δpyocins*, or 9 strong hits. Three biological replicates were performed with two technical replicates for each.

**Figure S6. R pyocins produced by PA14 and ST111 strains is bactericidal.** (**A**) Growth of S pyocin indicator strain PML1516d incubated with filtrates from self, ST253/PA14, ST253/PA14*Δpyocins*, or ST111 strains (ST111/M0067, ST111/M0101, ST111/218M0087). (**B**, **C**) Growth of ST291/M0103 (**B**) or ST299/M0128 (**C**) incubated with filtrates from self, ST111/218M0087, or ST111/218M0087*ΔRpyocin*. (**D**) CFUs (colony-forming unites) of ST260/M0089, ST291/M0103, and ST299/M0128 after 1 hour incubation with self, ST253/PA14, and ST253/PA14*Δpyocins* filtrates respectively. Three biological replicates were performed. Detection limit of CFU assay (horizontal dashed line) is 100 colony/mL per individual technical replicate.

**Figure S7. ST111 R pyocin not prophage contributes to the killing effect.** (**A, B**) Phage plaques formed on ST233/M0104 lawn with the treatment of 10 μL (**A**) or 200 μL (**B**) of filtrates from self, ST253/PA14, ST111/218M0087, or ST111/218M0087*ΔRpyocin*. Three biological replicates with three technical replicates each were performed. Scale bar: 0.5 cm.

**Figure S8. Deficiency in LPS biosynthesis is associated with the high susceptibility to R pyocins.** (**A**) The original pictures showing the swimming motility of *P. aeruginosa* PA14 mutants. A representative technical replicate is shown for each strain. The cropped pictures are shown in Fig 5A. (**B**) Growth of PA14 mutants incubated with filtrates from a R1 pyocin-producing strain ST260/M0089, a R2 pyocin-producing strain ST549/PAO1, or a R5 pyocin-producing strain ST111/218M0087 respectively. Three biological replicates with two technical replicates each were performed. (**C**) CFUs (colony-forming units) of the second line of PA14*ΔwaaL* after 1 hour incubation with filtrates from a R1 pyocin-producing strain ST260/M0089, a R2 pyocin-producing strain ST549/PAO1, or a R5 pyocin-producing strain ST111/218M0087 respectively. Three biological replicates were performed. Detection limit of CFU assay (horizontal dashed line) is 100 colony/mL per individual technical replicate.

**Figure S9. *waaL* deletion expands the killing range of R pyocins.** (**A**) CFUs (colony-forming units) of PA14*Δpyocins* and PA14*ΔpyocinsΔwaaL* (the second line) after 1 hour incubation with filtrates from self, a R1 pyocin-producing strain ST260/M0089, a R2 pyocin-producing strain ST549/PAO1, or a R5 pyocin**-**producing strain ST111/218M0087 respectively. (**B**) CFUs (colony-forming units) of 218M0087*ΔRpyocin* and 218M0087*ΔRpyocinΔwaaL* (the second line) after 1 hour incubation with filtrates from self, a R1 pyocin-producing strain ST260/M0089, a R2 pyocin**-** producing strain ST549/PAO1, or a R5 pyocin-producing strain ST111/218M0087 respectively. For (**A,B**), three biological replicates were performed. Detection limit of CFU assay (horizontal dashed line) is 100 colony/mL per individual technical replicate.

**Figure S10. The alignment of R pyocin fiber protein sequences from different subtypes.** (**A**) The alignment result showed the difference in R pyocin tail fiber protein sequences between R1 (ST260/M0089), R2 (ST253/PA14), R5 (ST111/218M0087). The color darkness indicates the identical amino acid residues between three sequences. (**B**) The phylogenetic tree based on the alignment of R pyocin fiber protein sequences from clinical isolates with different subtypes.

**Figure S11. MLST analysis of R1-4 pyocin-coding strains.** (**A, B**) Pie chart of MLST distribution of R1 pyocin-(**A**) or R2-4 pyocin-encoding strains (**B**). Strain numbers for each chart: 1,426 of R1 pyocin carriers and 1,202 of R2-4 pyocin encoding strains.

**Table S1. Growth inhibition by filtrates from isolates encoding different R pyocins**

## Data Availability Statement

The authors confirm that the data supporting the findings of this study are available within the article and its supplementary materials.

## Author Contributions

NK contributed to the conception and design of the study. LZ, FT, QX and YD conducted experiments, organized the database, and performed the statistical analysis. MH and SS conducted epidemiological study and collected clinical isolates. LZ, FT, QX, and NK drafted the manuscript. NK funded the research and provided overall supervision of the project. All authors contributed to manuscript revision, read, and approved the submitted version.

