## Supplementary material for "Role of R5 Pyocin in the Predominance of High-Risk *Pseudomonas aeruginosa* Isolates": Combined Sup Materials

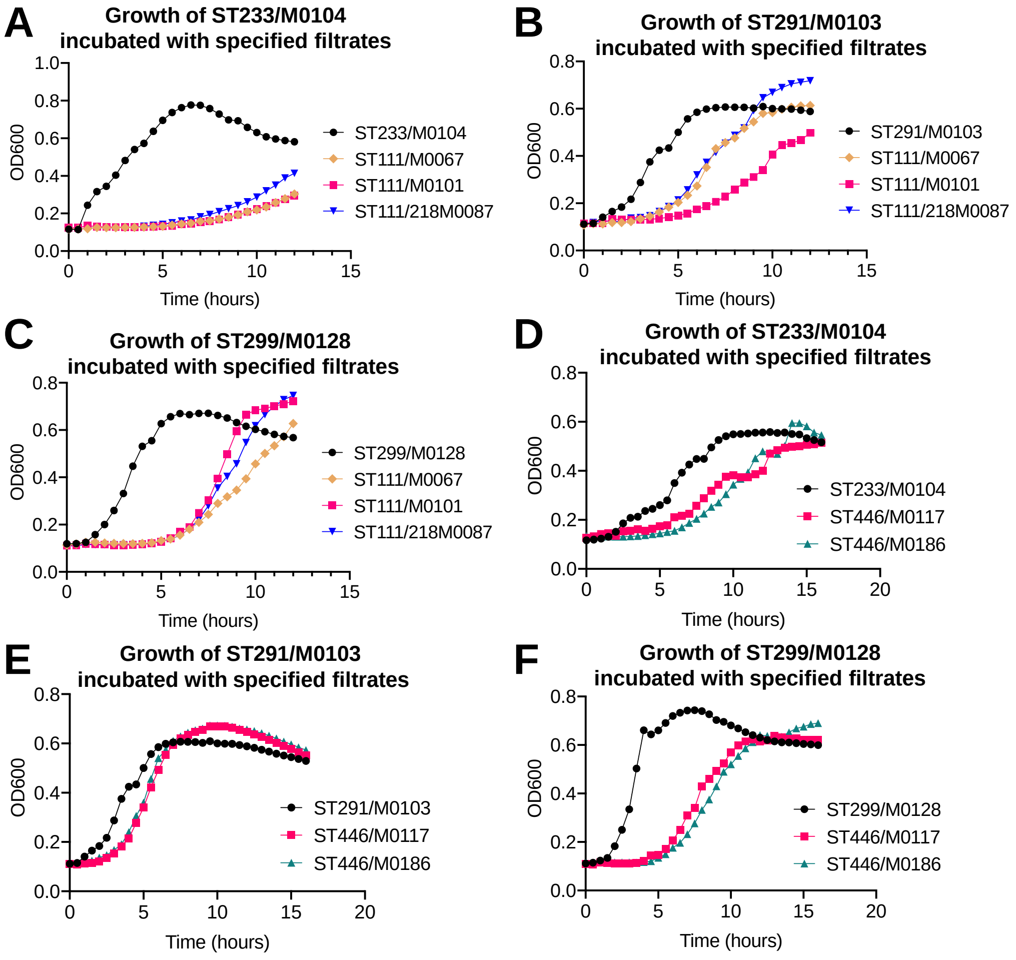
A
Growth of ST233/M0104
incubated with specified filtrates
OD600
Time (hours)
ST233/M0104
ST111/M0067
ST111/M0101
ST111/218M0087
B
Growth of ST291/M0103
incubated with specified filtrates
OD600
Time (hours)
ST291/M0103
ST111/M0067
ST111/M0101
ST111/218M0087
C
Growth of ST299/M0128
incubated with specified filtrates
OD600
Time (hours)
ST299/M0128
ST111/M0067
ST111/M0101
ST111/218M0087
D
Growth of ST233/M0104
incubated with specified filtrates
OD600
Time (hours)
ST233/M0104
ST446/M0117
ST446/M0186
E
Growth of ST291/M0103
incubated with specified filtrates
OD600
Time (hours)
ST291/M0103
ST446/M0117
ST446/M0186
F
Growth of ST299/M0128
incubated with specified filtrates
OD600
Time (hours)
ST299/M0128
ST446/M0117
ST446/M0186

### Figure S1

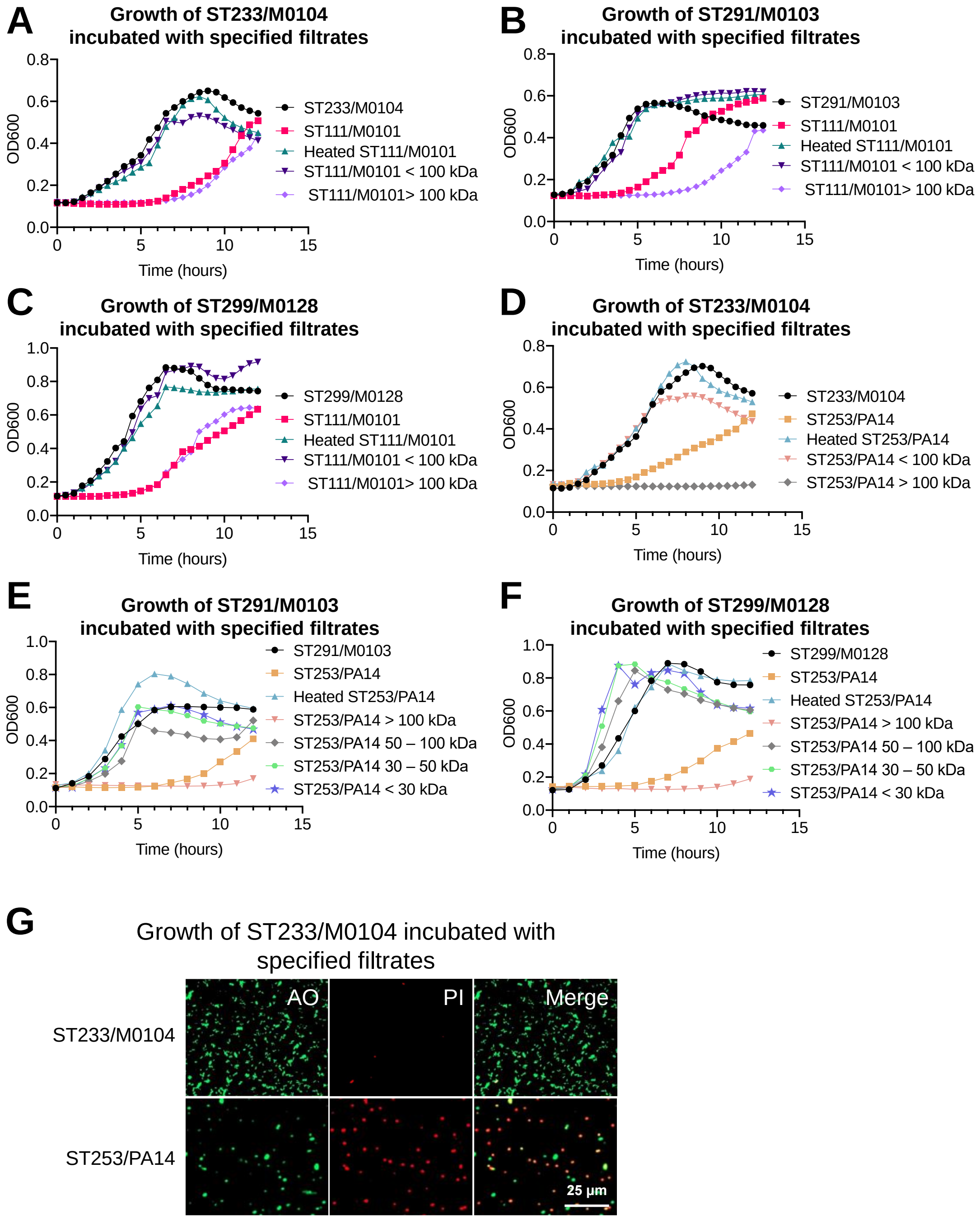
A
Growth of ST233/M0104
incubated with specified filtrates
OD600
Time (hours)
ST233/M0104
ST111/M0101
Heated ST111/M0101
ST111/M0101 < 100 kDa
ST111/M0101> 100 kDa
B
Growth of ST291/M0103
incubated with specified filtrates
OD600
Time (hours)
ST291/M0103
ST111/M0101
Heated ST111/M0101
ST111/M0101 < 100 kDa
ST111/M0101> 100 kDa
C
Growth of ST299/M0128
incubated with specified filtrates
OD600
Time (hours)
ST299/M0128
ST111/M0101
Heated ST111/M0101
ST111/M0101 < 100 kDa
ST111/M0101> 100 kDa
D
Growth of ST233/M0104
incubated with specified filtrates
OD600
Time (hours)
ST233/M0104
ST253/PA14
Heated ST253/PA14
ST253/PA14 < 100 kDa
ST253/PA14 > 100 kDa
E
Growth of ST291/M0103
incubated with specified filtrates
OD600
Time (hours)
ST291/M0103
ST253/PA14
Heated ST253/PA14
ST253/PA14 > 100 kDa
ST253/PA14 50 – 100 kDa
ST253/PA14 30 – 50 kDa
ST253/PA14 < 30 kDa
F
Growth of ST299/M0128
incubated with specified filtrates
OD600
Time (hours)
ST299/M0128
ST253/PA14
Heated ST253/PA14
ST253/PA14 > 100 kDa
ST253/PA14 50 – 100 kDa
ST253/PA14 30 – 50 kDa
ST253/PA14 < 30 kDa
G
Growth of ST233/M0104 incubated with specified filtrates
AO
PI
Merge
ST233/M0104
ST253/PA14
25 µm

**Figure S2**

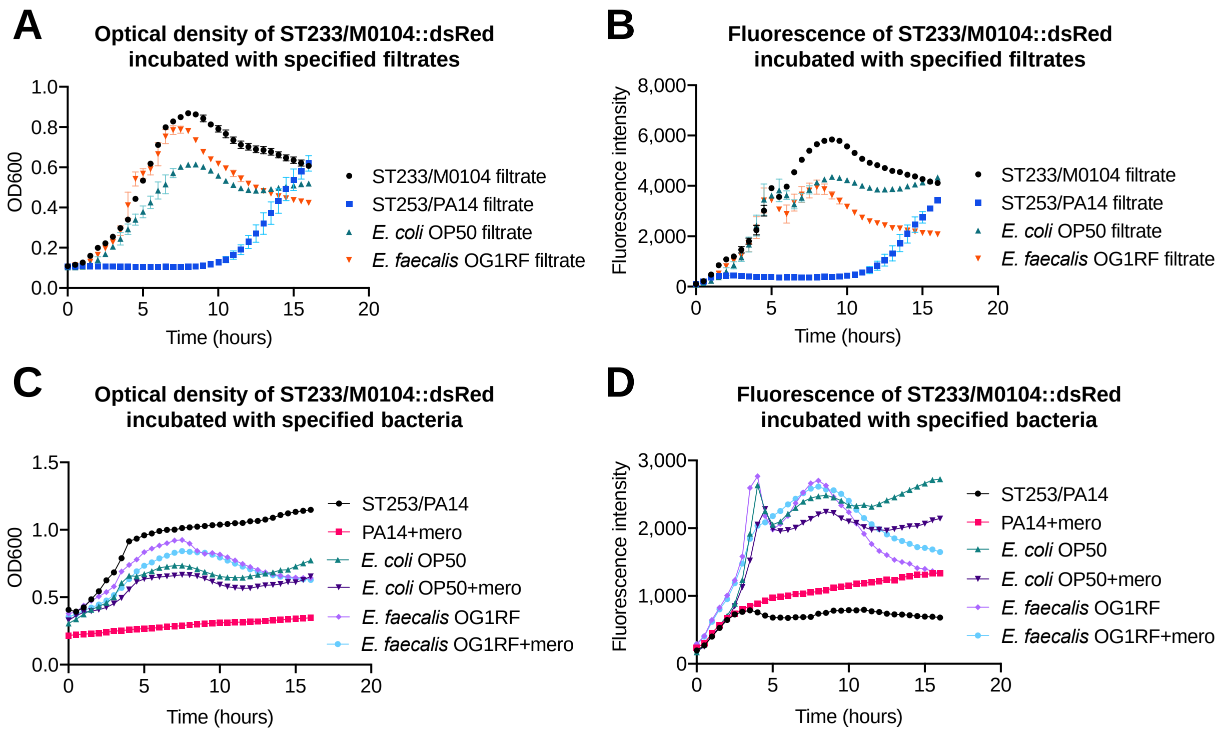
A
Optical density of ST233/M0104::dsRed incubated with specified filtrates
OD600
Time (hours)
ST233/M0104 filtrate
ST253/PA14 filtrate
E. coli OP50 filtrate
E. faecalis OG1RF filtrate
B
Fluorescence of ST233/M0104::dsRed incubated with specified filtrates
Fluorescence intensity
Time (hours)
ST233/M0104 filtrate
ST253/PA14 filtrate
E. coli OP50 filtrate
E. faecalis OG1RF filtrate
C
Optical density of ST233/M0104::dsRed incubated with specified bacteria
OD600
Time (hours)
ST253/PA14
PA14+mero
E. coli OP50
E. coli OP50+mero
E. faecalis OG1RF
E. faecalis OG1RF+mero
D
Fluorescence of ST233/M0104::dsRed incubated with specified bacteria
Fluorescence intensity
Time (hours)
ST253/PA14
PA14+mero
E. coli OP50
E. coli OP50+mero
E. faecalis OG1RF
E. faecalis OG1RF+mero

### Figure S3

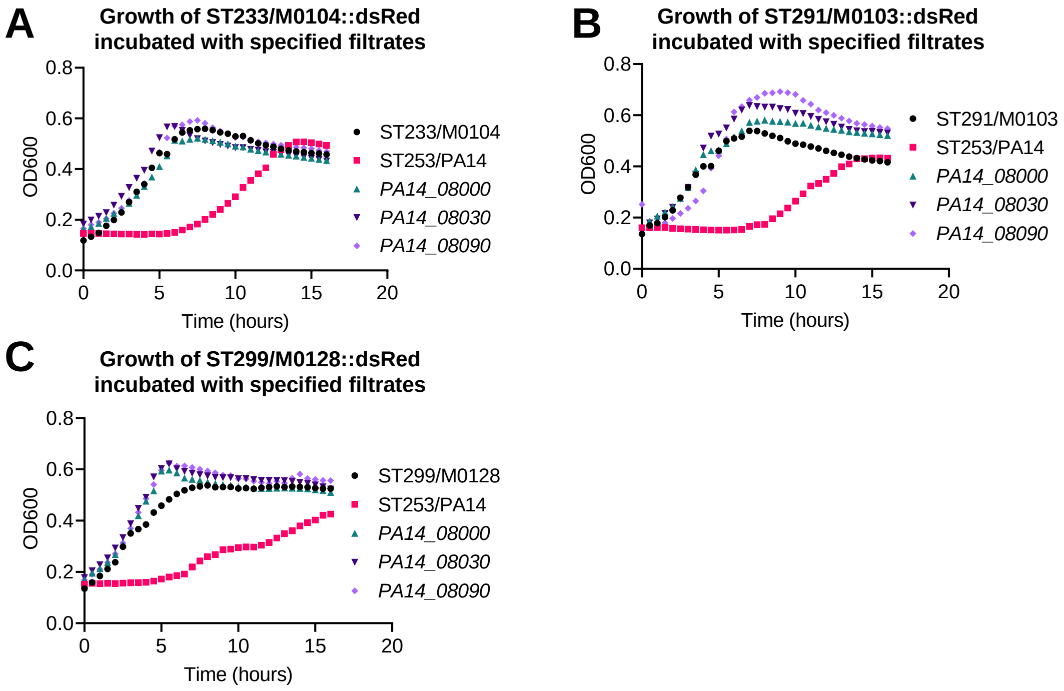

A
Growth of ST233/M0104::dsRed
incubated with specified filtrates
OD600
Time (hours)
ST233/M0104
ST253/PA14
PA14_08000
PA14_08030
PA14_08090
B
Growth of ST291/M0103::dsRed
incubated with specified filtrates
OD600
Time (hours)
ST291/M0103
ST253/PA14
PA14_08000
PA14_08030
PA14_08090
C
Growth of ST299/M0128::dsRed
incubated with specified filtrates
OD600
Time (hours)
ST299/M0128
ST253/PA14
PA14_08000
PA14_08030
PA14_08090


### Figure S4

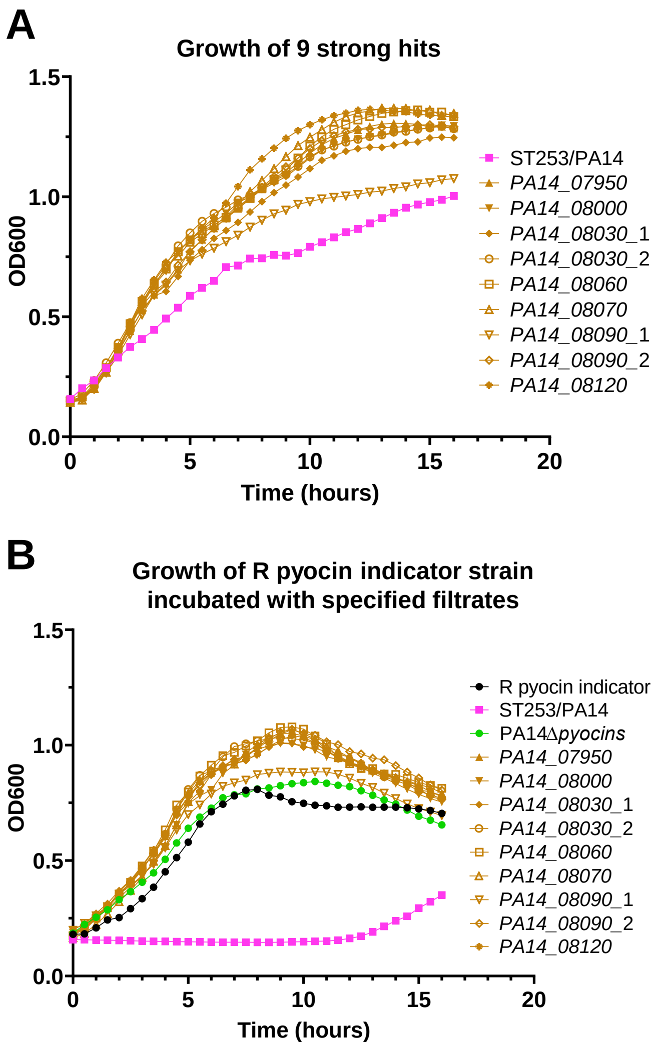
A
Growth of 9 strong hits
OD600
Time (hours)
0.0
0.5
1.0
1.5
0
5
10
15
20
ST253/PA14
PA14_07950
PA14_08000
PA14_08030_1
PA14_08030_2
PA14_08060
PA14_08070
PA14_08090_1
PA14_08090_2
PA14_08120
B
Growth of R pyocin indicator strain
incubated with specified filtrates
OD600
Time (hours)
0.0
0.5
1.0
1.5
0
5
10
15
20
R pyocin indicator
ST253/PA14
PA14Δpyocins
PA14_07950
PA14_08000
PA14_08030_1
PA14_08030_2
PA14_08060
PA14_08070
PA14_08090_1
PA14_08090_2
PA14_08120

### Figure S5

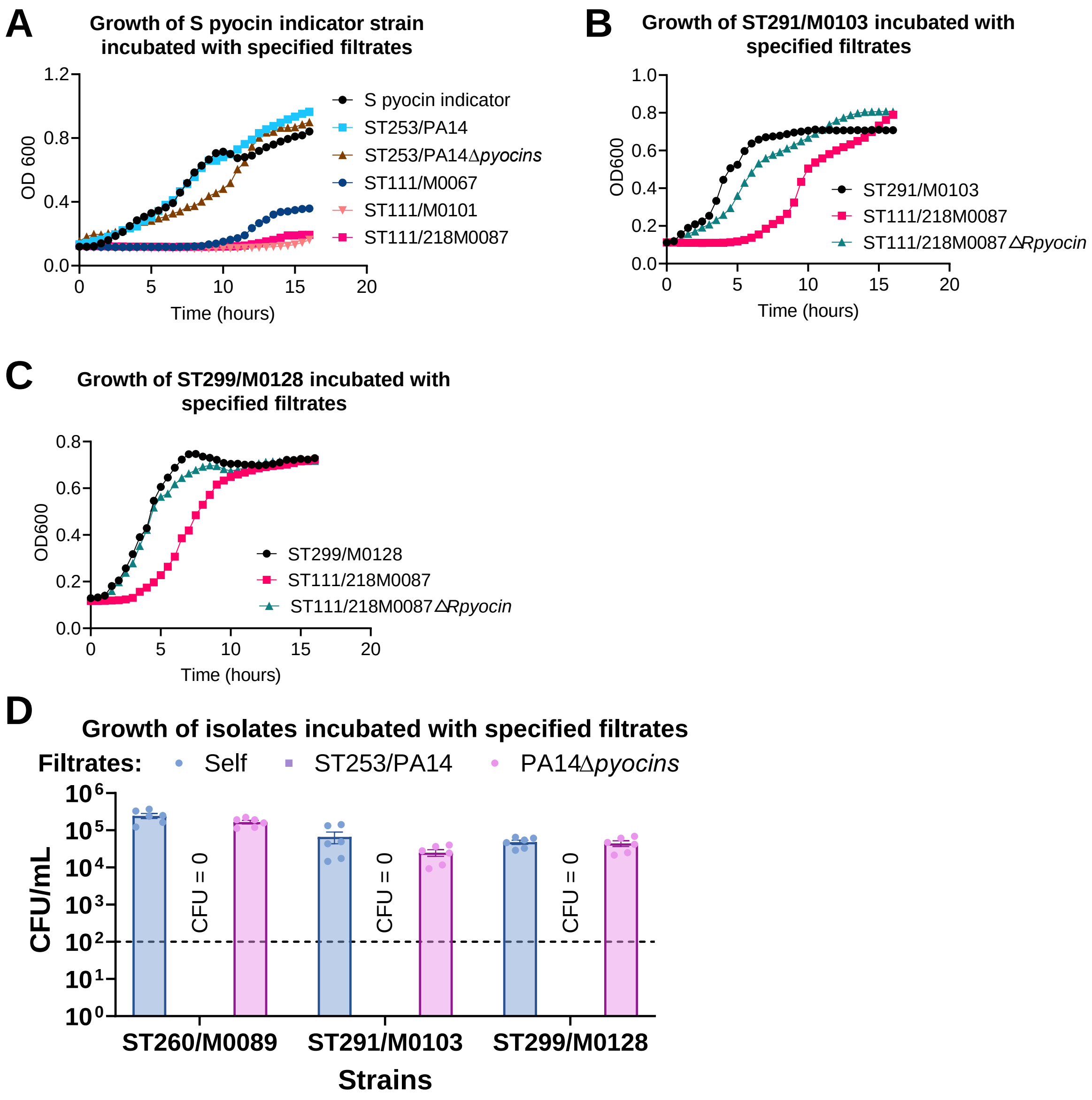

A
Growth of S pyocin indicator strain incubated with specified filtrates
OD 600
Time (hours)
S pyocin indicator
ST253/PA14
ST253/PA14Δpyocins
ST111/M0067
ST111/M0101
ST111/218M0087
B
Growth of ST291/M0103 incubated with specified filtrates
OD600
Time (hours)
ST291/M0103
ST111/218M0087
ST111/218M0087ΔRpyocin
C
Growth of ST299/M0128 incubated with specified filtrates
OD600
Time (hours)
ST299/M0128
ST111/218M0087
ST111/218M0087ΔRpyocin
D
Growth of isolates incubated with specified filtrates
Filtrates: Self ST253/PA14 PA14Δpyocins
CFU/mL
CFU = 0
ST260/M0089
ST291/M0103
ST299/M0128
Strains


**Figure S6**

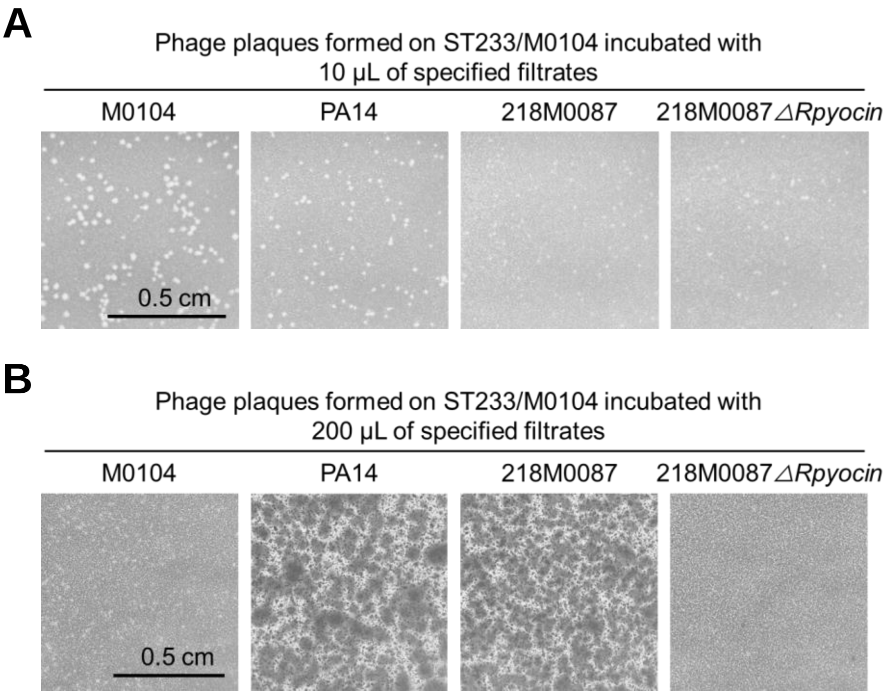
A
Phage plaques formed on ST233/M0104 incubated with
10 μL of specified filtrates
M0104
PA14
218M0087
218M0087△Rpyocin
0.5 cm
B
Phage plaques formed on ST233/M0104 incubated with
200 μL of specified filtrates
M0104
PA14
218M0087
218M0087△Rpyocin
0.5 cm

**Figure S7**

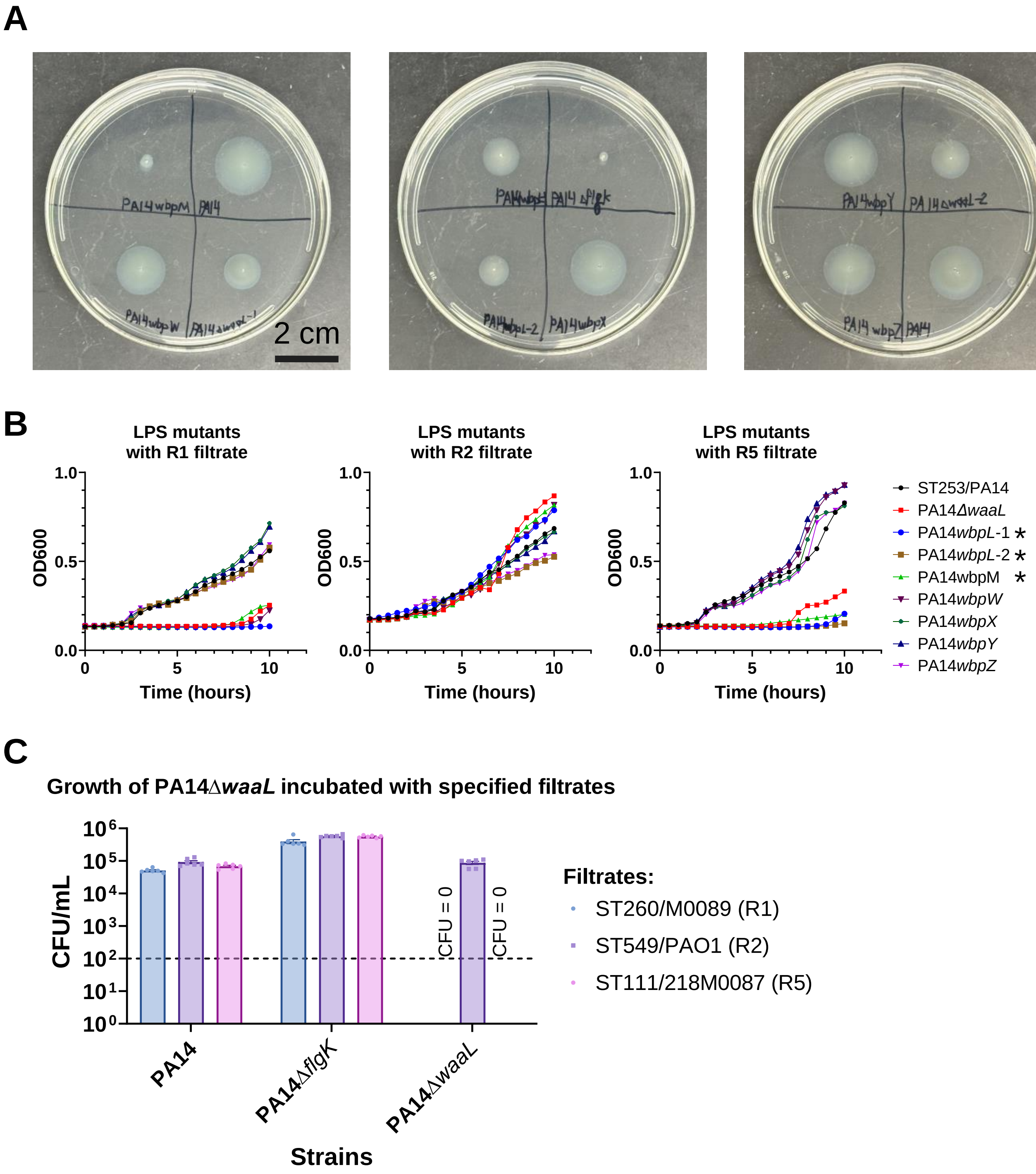
A
PA14wbpM
PA14
PA14wbpW
PA14ΔwaaL-1
2 cm
PA14wbpL-2
PA14wbpX
PA14wbpY
PA14ΔwaaL-2
PA14 wbpZ
PA14
B
LPS mutants with R1 filtrate
LPS mutants with R2 filtrate
LPS mutants with R5 filtrate
OD600
Time (hours)
ST253/PA14
PA14ΔwaaL
PA14wbpL-1 *
PA14wbpL-2 *
PA14wbpM *
PA14wbpW
PA14wbpX
PA14wbpY
PA14wbpZ
C
Growth of PA14ΔwaaL incubated with specified filtrates
CFU/mL
CFU = 0
CFU = 0
PA14
PA14ΔflgK
PA14ΔwaaL
Strains
Filtrates:
ST260/M0089 (R1)
ST549/PAO1 (R2)
ST111/218M0087 (R5)

### Figure S8

**A**

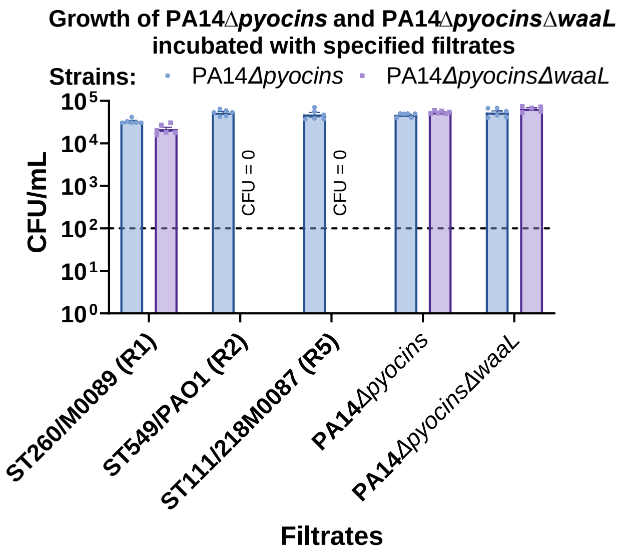
Growth of PA14ΔpyocinsΔ and PA14ΔpyocinsΔwaaL
incubated with specified filtrates
Strains: PA14Δpyocins PA14ΔpyocinsΔwaaL
CFU/mL
10^5
10^4
10^3
10^2
10^1
10^0
CFU = 0
CFU = 0
ST260/M0089 (R1)
ST549/PAO1 (R2)
ST111/218M0087 (R5)
PA14Δpyocins
PA14ΔpyocinsΔwaaL
Filtrates

**B**

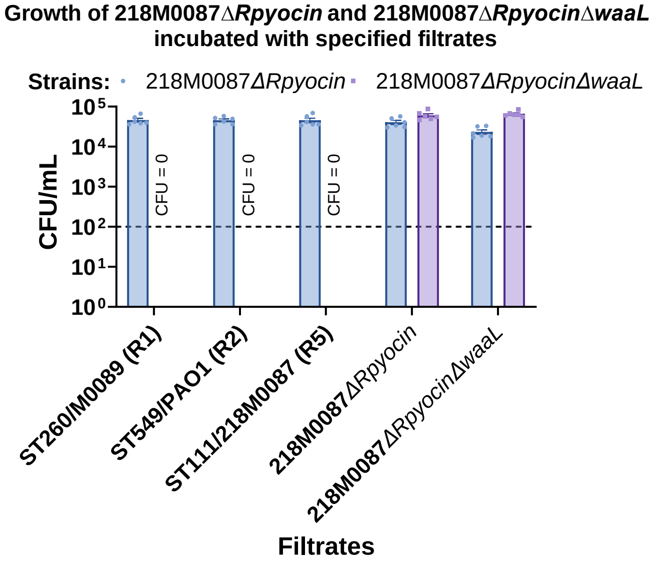
Growth of 218M0087ΔRpyocin and 218M0087ΔRpyocinΔwaaL
incubated with specified filtrates
Strains: 218M0087ΔRpyocin 218M0087ΔRpyocinΔwaaL
CFU/mL
10^5
10^4
10^3
10^2
10^1
10^0
CFU = 0
CFU = 0
CFU = 0
ST260/M0089 (R1)
ST549/PAO1 (R2)
ST111/218M0087 (R5)
218M0087ΔRpyocin
218M0087ΔRpyocinΔwaaL
Filtrates

### Figure S9

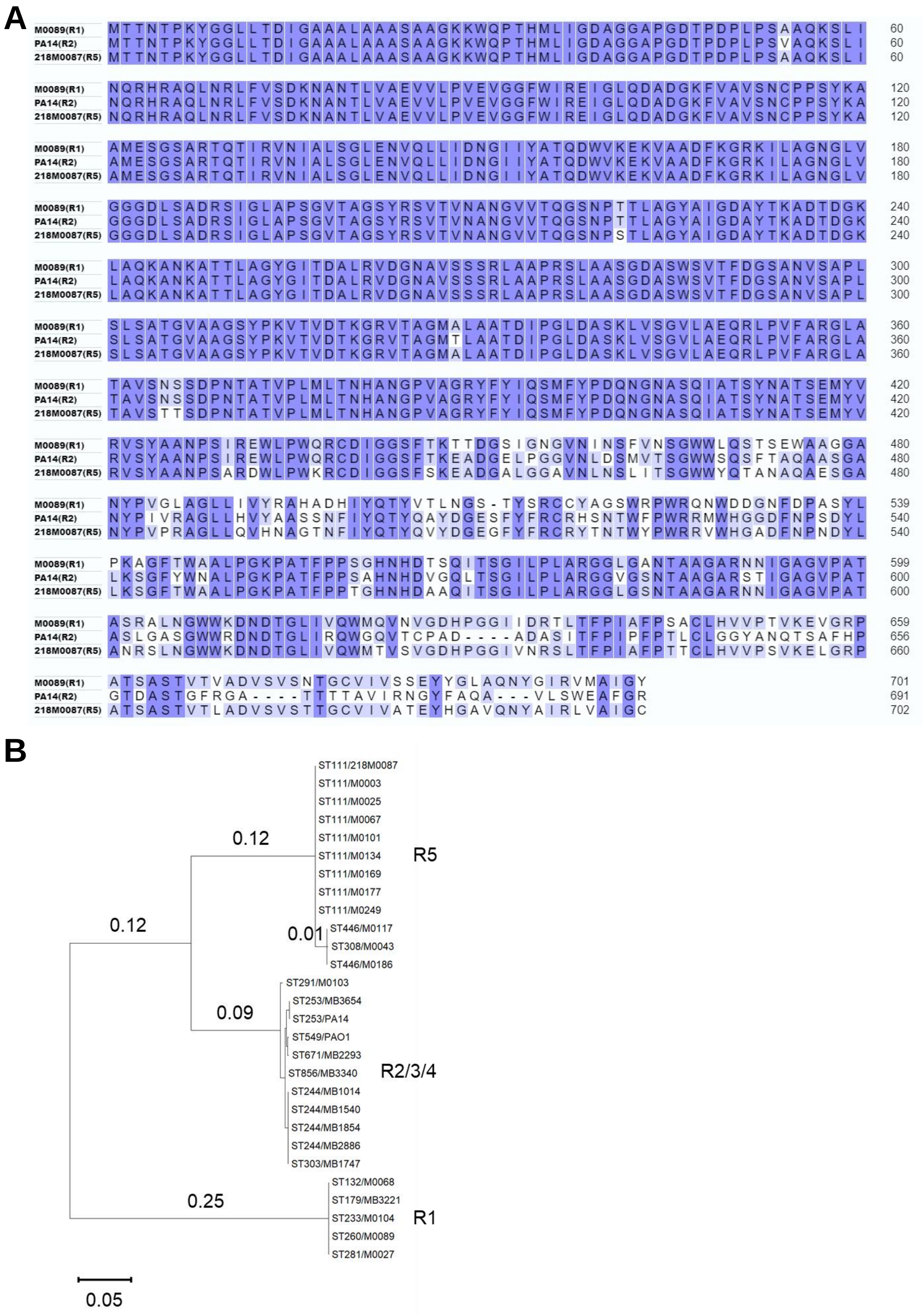
A
M0089(R1) MTTNTPKYGGLLTDIGAAALAAASAAGKKWQPTHMLIGDAGGAPGDTPDPLPSAAQKSLI 60
PA14(R2) MTTNTPKYGGLLTDIGAAALAAASAAGKKWQPTHMLIGDAGGAPGDTPDPLPSVAQKSLI 60
218M0087(R5) MTTNTPKYGGLLTDIGAAALAAASAAGKKWQPTHMLIGDAGGAPGDTPDPLPSAAQKSLI 60
M0089(R1) NQRHRAQLNRLFVSDKNANTLVAEVVLPVEVGGFWIREIGLQDADGKFVAVSNCPPSYKA 120
PA14(R2) NQRHRAQLNRLFVSDKNANTLVAEVVLPVEVGGFWIREIGLQDADGKFVAVSNCPPSYKA 120
218M0087(R5) NQRHRAQLNRLFVSDKNANTLVAEVVLPVEVGGFWIREIGLQDADGKFVAVSNCPPSYKA 120
M0089(R1) AMESGSARTQTIRVNIALSGLENVQLLIDNGIIYATQDWVKEKVAADFKGRKILAGNGLV 180
PA14(R2) AMESGSARTQTIRVNIALSGLENVQLLIDNGIIYATQDWVKEKVAADFKGRKILAGNGLV 180
218M0087(R5) AMESGSARTQTIRVNIALSGLENVQLLIDNGIIYATQDWVKEKVAADFKGRKILAGNGLV 180
M0089(R1) GGGDLSADRSIGLAPSGVTAGSYRSVTVNANGVVTQGSNPTTLAGYAIGDAYTKADTDGK 240
PA14(R2) GGGDLSADRSIGLAPSGVTAGSYRSVTVNANGVVTQGSNPTTLAGYAIGDAYTKADTDGK 240
218M0087(R5) GGGDLSADRSIGLAPSGVTAGSYRSVTVNANGVVTQGSNPSTLAGYAIGDAYTKADTDGK 240
M0089(R1) LAQKANKATTLAGYGITDALRVDGNAVSSSRLAAPRSLAASGDASWSVTFDGSANVSAPL 300
PA14(R2) LAQKANKATTLAGYGITDALRVDGNAVSSSRLAAPRSLAASGDASWSVTFDGSANVSAPL 300
218M0087(R5) LAQKANKATTLAGYGITDALRVDGNAVSSSRLAAPRSLAASGDASWSVTFDGSANVSAPL 300
M0089(R1) SLSATGVAAGSYPKVTVDTKGRVTAGMALAATDIPGLDASKLVSGVLAEQRLPVFARGLA 360
PA14(R2) SLSATGVAAGSYPKVTVDTKGRVTAGMTLAATDIPGLDASKLVSGVLAEQRLPVFARGLA 360
218M0087(R5) SLSATGVAAGSYPKVTVDTKGRVTAGMALAATDIPGLDASKLVSGVLAEQRLPVFARGLA 360
M0089(R1) TAVSNSSDPNTATVPLMLTNHANGPVAGRYFYIQSMFYPDQNGNASQIATSYNATSEMYV 420
PA14(R2) TAVSNSSDPNTATVPLMLTNHANGPVAGRYFYIQSMFYPDQNGNASQIATSYNATSEMYV 420
218M0087(R5) TAVSTTSDPNTATVPLMLTNHANGPVAGRYFYIQSMFYPDQNGNASQIATSYNATSEMYV 420
M0089(R1) RVSYAANPSIREWLPWQRCDIGGSFTKTTDGSIGNGVNINSFVNSGWWLQSTSEWAAGGA 480
PA14(R2) RVSYAANPSIREWLPWQRCDIGGSFTKEADGELPGGVNLDSMVTSGWWSQSFTAQAASGA 480
218M0087(R5) RVSYAANPSARDWLPWKRCDIGGSFSKEADGALGGAVNLNSLITSGWWYQTANAQAESGA 480
M0089(R1) NYPVGLAGLLIVYRAHADHIYQTYVTLNGS-TYSRCCYAGSWRPWRQNWDDGNFDPASYL 539
PA14(R2) NYPIVRAGLLHVYAASSNFIYQTYQAYDGESFYFRCRHSNTWFPWRRMWHGGDFNPSDYL 540
218M0087(R5) NYPVPRAGLLQVHNAGTNFIYQTYQVYDGEGFYFRCRYTNTWYPWRRVWHGADFNPNDYL 540
M0089(R1) PKAGFTWAALPGKPATFPPSGHNHDTSQITSGILPLARGGLGANTAAGARNNIGAGVPAT 599
PA14(R2) LKSGFYWNALPGKPATFPPSAHNHDVGQLTSGILPLARGGVGSNTAAGARSTIGAGVPAT 600
218M0087(R5) LKSGFTWAALPGKPATFPPTGHNHDAAQITSGILPLARGGLGSNTAAGARNNIGAGVPAT 600
M0089(R1) ASRALNGWWKDNDTGLIVQWMQVNVGDHPGGIIDRTLTFPIAFPSACLHVVPTVKEVGRP 659
PA14(R2) ASLGASGWWRDNDTGLIRQWGQVTCPAD----ADASITFPIPFPTLCLGGYANQTSAFHP 656
218M0087(R5) ANRSLNGWWKDNDTGLIVQWMTVSVGDHPGGIVNRSLTFPIAFPTTCLHVVPSVKELGRP 660
M0089(R1) ATSASTVTVADVSVSNTGCVIVSSEYYGLAQNYGIRVMAIGY 701
PA14(R2) GTDASTGFRGA----TTTTAVIRNGYFAQA---VLSWEAFGR 691
218M0087(R5) ATSASTVTLADVSVSTTGCVIVATEYHGAVQNYAIRLVAIGC 702
B
ST111/218M0087
ST111/M0003
ST111/M0025
ST111/M0067
ST111/M0101
ST111/M0134
ST111/M0169
ST111/M0177
ST111/M0249
ST446/M0117
ST308/M0043
ST446/M0186
R5
0.12
0.12
0.01
ST291/M0103
ST253/MB3654
ST253/PA14
ST549/PAO1
ST671/MB2293
ST856/MB3340
ST244/MB1014
ST244/MB1540
ST244/MB1854
ST244/MB2886
ST303/MB1747
R2/3/4
0.09
ST132/M0068
ST179/MB3221
ST233/M0104
ST260/M0089
ST281/M0027
R1
0.25
0.05

### Figure S10

A

MLST distribution of R1 pyocin encoding strains

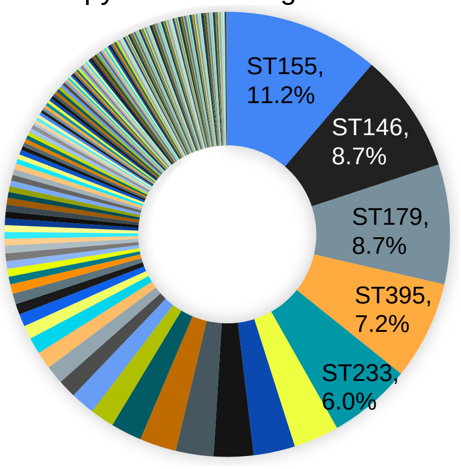


Total strains: 1,426
(of which 0 ST235, 0 ST111, and 0 ST446)

B

MLST distribution of R2-4 pyocin encoding strains

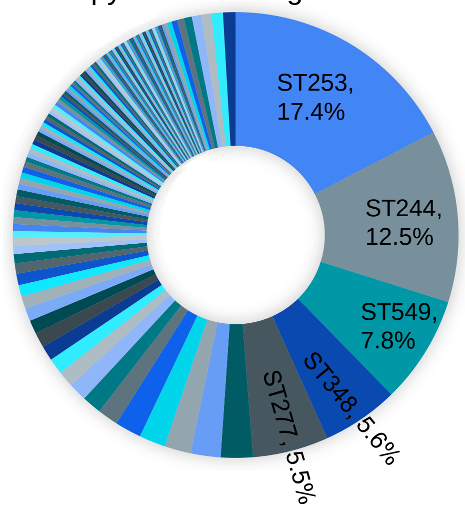


Total strains: 1,202
(of which 0 ST235, 0 ST111, and 0 ST446)

**Figure S11**

#### Table S1. Growth inhibition by filtrates from isolates encoding different R pyocins

| Inhibit the growth of | MLST | M0027 (R1) 281 | M0089 (R1) 260 | PA14 (R2) 253 | PAO1 (R2) 549 | M0067 (R5) 111 | M0101 (R5) 111 | 218M0087 (R5) 111 |
|---|---|---|---|---|---|---|---|---|
| M0067 (R5) | 111 | − | − | − | − | − | − | − |
| M0101 (R5) | 111 | − | − | − | − | − | − | − |
| 218M0087 (R5) | 111 | − | − | − | − | − | − | − |
| M0003 (R5) | 111 | + | + | − | − | − | − | − |
| M0025 (R5) | 111 | − | − | − | − | − | − | − |
| M0134 (R5) | 111 | − | − | − | − | − | − | − |
| M0169 (R5) | 111 | − | − | − | − | − | − | − |
| M0177 (R5) | 111 | − | − | − | − | − | − | − |
| M0249 (R5) | 111 | − | − | − | − | − | − | − |
| M0117 (R5) | 446 | + | + | − | − | − | − | − |
| M0186 (R5) | 446 | − | − | − | − | − | − | − |
| M0068 (R1) | 132 | − | − | + | + | + | + | + |
| M0103 (R2) | 291 | + | + | + | + | + | + | + |
| M0104 (R1) | 233 | + | − | + | + | + | + | + |
| M0128 (No Match) | 299 | + | + | + | + | + | + | + |
| M0013 (No Match) | 17 | + | + | + | + | + | + | + |
| M0027 (R1) | 281 | − | − | + | + | + | + | + |
| M0043 (R5) | 308 | − | + | + | + | + | + | + |
| M0089 (R1) | 260 | − | − | + | + | + | + | + |

MLST: Multilocus sequence typing; **−**: not inhibit; **+**: inhibit
